# Assessing the inference of single-cell phylogenies and population dynamics from CRISPR lineage recordings

**DOI:** 10.1101/2025.01.08.631854

**Authors:** Julia Pilarski, Tanja Stadler, Sophie Seidel

## Abstract

Multicellular organisms develop from a single cell by repeated rounds of cell division, differentiation, and death, which can be represented as a single-cell phylogenetic tree. Genetic lineage tracing allows us to investigate this development by tracking the ancestry of individual cells as populations grow and change over time. However, accurate reconstruction of the cell phylogeny and quantification of the corresponding phylodynamic parameters - cell division, differentiation and death rates - from this tracking data remains challenging and needs to be systematically evaluated.

We perform simulations and assess, using the Bayesian framework, the joint inference of time-scaled cell phylogenies and phylodynamic parameters from CRISPR lineage recordings with random or sequential edits. Principally, we characterize the inference improvements as the recorder capacity increases. We observe more accurate phylogenetic reconstruction from sequential compared to random recordings, but no substantial improvement in phylodynamic inference when using the additional information contained in the order of edits. Overall, we find that CRISPR lineage recordings carry a strong signal on the rates of cell division when appropriate models are used. However, we detect biases in the inferred rates of cell division and death under phylodynamic model misspecification, i.e. when fitting classic memoryless birth-death processes to synchronous cell divisions.

Moreover, for scenarios when cells differentiate into distinct types, we demonstrate that Bayesian phylodynamic analysis of sparse end-point measurements can resolve these cell differentiation trajectories by lineage and time. Under prototypical dynamics, we recover cell type-specific division and death rates, and cell type transition rates in over **80%** of simulations.

Overall, this simulation study explores how much information on cellular development can be extracted from state-of-the-art genetic lineage tracing data using phylogenetic and phylodynamic methodology.

**Author summary:** Novel technologies provide means to trace the development of cell populations over time by introducing heritable and editable genetic sequences that record lineage information in the cells’ genome. Reconstructing a population’s history from such sequences sparsely sampled at a single time point is, however, computationally challenging. In this work, we use simulations and statistical inference to evaluate how accurately we can recover the relationships among cells and estimate the temporal dynamics of cell populations from genetic lineage tracing data generated from distinct recording systems, and compare their information content. Our results show that it is possible to quantify how cells divide, differentiate, and die based on such data, though certain statistical limitations remain. Addressing these limitations in future research will be essential for deepening our understanding of cell development in complex tissues and organisms, in both health and disease.

## 1 Introduction

Uncovering how cells proliferate and specialize is crucial for understanding fundamental biological processes such as the development of multicellular organisms from a single cell, tissue regeneration, and disease progression. In recent years, genetic lineage tracing has emerged as a powerful technology for recording the lineages of individual cells over time. Increasingly, CRISPR-Cas systems are leveraged to induce heritable and irreversible mutations (‘edits’) in short DNA sequences (‘targets’ or ‘barcodes’) which are introduced into the cells’ genomes [1–5]. Targets accumulate edits over time as they are passed on from cells to their descendants. At the end of this process, the diverse barcodes can be retrieved by single-cell sequencing and used to reconstruct cell lineage trees (‘phylogenetic trees’ or ‘phylogenies’). Such trees, in turn, contain information on how cell populations grow and differentiate, and can be analyzed within the statistical framework of phylodynamics [6].

Several CRISPR lineage recorders have been developed in the past decade. Many of them consist of multiple targets, each of which is a CRISPR-Cas9 target site that independently acquires indel mutations upon activation of the editing reagents [7–10] (‘non-sequential’ recorders). Novel tools employ prime editors that introduce short template-based insertions in tandem arrays (‘tapes’) of target sites, enabling an ordered recording of edits [11–13] (‘sequential’ recorders). These recently emerging prime editing-based lineage recorders aim to resolve individual cell divisions and trace cell lineages across temporal scales.

Despite technological advancements, it remains computationally challenging to accurately infer the cell relationships from mutations observed in a sample of cells at a single point in time [1]. Previous studies [14–16] have demonstrated limits of phylogeny reconstruction from CRISPR lineage recordings. In particular, they have shown that constraints on recorder capacity, the editing rate, and diversity of editing outcomes can result in not completely resolved phylogenies. Nevertheless, the studies have identified, using simulation and theory, experimental conditions over which exact tree reconstruction is possible.

Importantly, non-sequential and sequential recorders have been investigated separately until now, and a comparison of their information content is lacking. In particular, it is not clear how much information sequential editing quantitatively adds to phylogenetic reconstruction. Moreover, little attention has been paid so far to the quantification of population dynamics, i.e., the dynamics of cell division, differentiation, and death, based on CRISPR lineage recordings. We aim to fill the gap by evaluating the inference of phylodynamic parameters, alongside cell phylogenies, from the data. Given the absence of ground truth information for most lineage tracing experiments, we perform simulations. We mimic lineage tracing experiments under a wide range of scenarios and simulate data for both non-sequential and sequential recorders. Then, we apply phylodynamic models and assess how much signal the data contains to accurately infer the cell phylogeny and the population dynamics. In particular, we employ the Bayesian framework, allowing us to incorporate prior knowledge into the analysis and capture the uncertainty associated with the inference results.

Bayesian phylogenetic reconstruction methods have been recently tailored to CRISPR lineage recordings [17–19], but phylodynamic models have not yet been thoroughly investigated in the context of cell biology. A central assumption in many phylodynamic models – particularly birth-death models, established and widely used in epidemiology and macroevolution [20] – is that branching (here, cell division) occurs according to a Poisson process, implying exponentially distributed waiting times and memoryless behavior. However, this assumption may not hold in cell development which, at least during early embryonic stages, is known to follow more synchronous and regular patterns [21–24].

Another key challenge is inferring differentiation maps from single-cell lineage tracing data, particularly when cells are sequenced only at a final time point. Although editable, heritable genetic barcodes track the lineage relationships arising through cell divisions, they do not directly measure transitions between cell types. Several recent approaches have sought to address this limitation by integrating lineage information with end-point cell state measurements [25–30]. However, these methods have focused on characterizing differentiation based on reconstructed cell lineage trees, rather than jointly inferring the trees and differentiation dynamics from sequence data using a stochastic branching process as the tree prior.

Here, we address the adequacy and applicability of phylodynamic birth-death processes to cell population dynamics. We first explore basic principles for homogeneous cell populations and then adopt the phylodynamic approach to infer population dynamics that vary across cell types. Further, we explore whether – and to what extent – past cell differentiation trajectories can be recovered along time-scaled lineage trees by coupling a multi-type phylodynamic birth-death model to the CRISPR editing models in the Bayesian framework.

In particular, our first goal is to compare the phylogenetic and phylodynamic signal carried by non-sequential and sequential CRISPR lineage recorders. By systematically altering the number of editable target sites, the editing rate, and the editing window in our simulations, we show that recorder design and capacity significantly affect the inference performance. Second, we investigate in what scenarios the rates of cell division and death can be estimated reliably from the recorder data. We identify important limitations of the currently available phylodynamic models to capture realistic cell population dynamics. Third, we aim to infer cell differentiation dynamics from simulated lineage recordings in which sampled cells are annotated with their cell types. In summary, we find that sparse end-point measurements contain rich information on cell type-specific division and death rates, and cell type transition rates.

## 2 Results

### 2.1 The workflow

To evaluate and compare the information contained in CRISPR lineage tracing data, we established the following workflow Fig. 1 (for details, see Methods): First, we simulated time-scaled cell phylogenies which represent the development of cell populations from a single ancestor. Note that we use here the terms ‘phylogeny’ and ‘phylogenetic tree’ interchangeably to refer to the ancestry of a sample of cells from a population. Others might call that object also ‘lineage tree’, as discussed in [6]. We considered multiple tree generating processes (population dynamics or phylodynamic models) and simulated 20 trees per model (see Fig. S2 for examples). Each process was motivated by realistic cell population dynamics, from completely synchronous divisions in a homogeneous cell population resembling early embryonic development, to stochastic divisions in a heterogeneous cell population resembling differentiating cells.

**Fig. 1:**
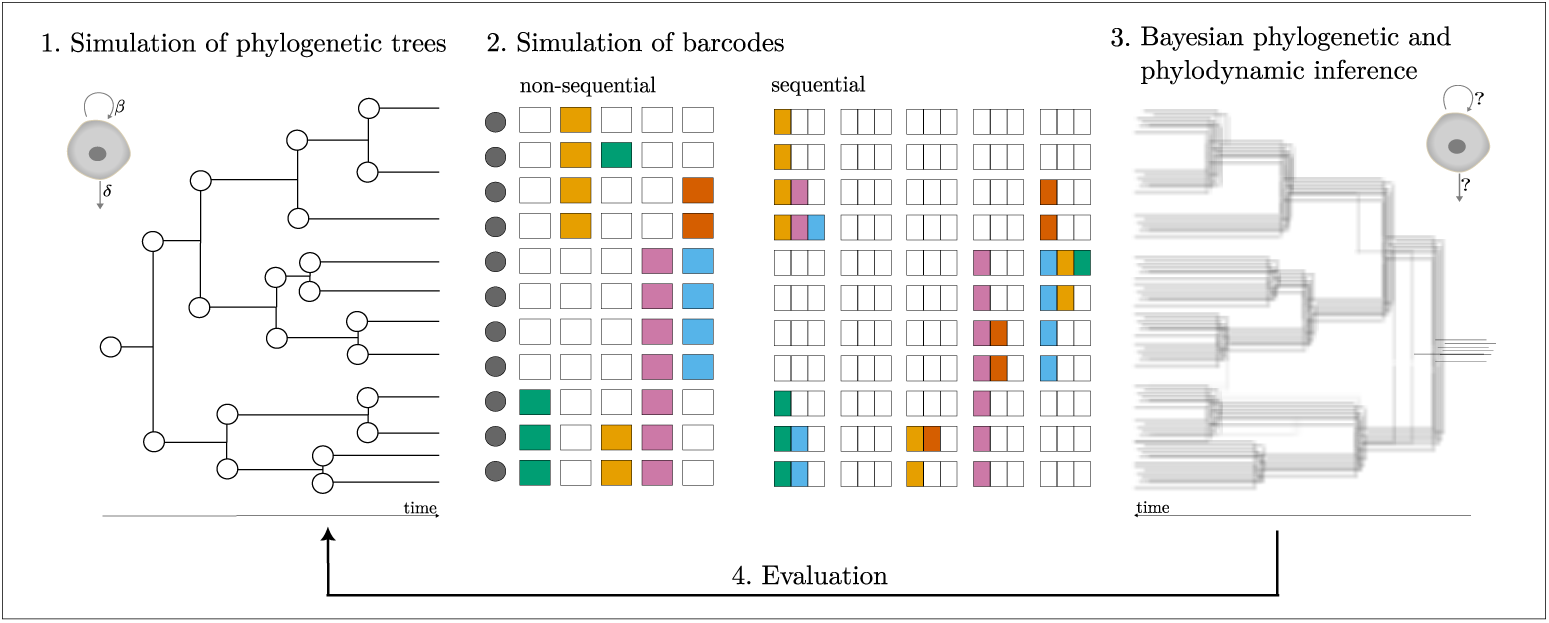
Schematic of the workflow. First, we generated phylogenetic trees according to different phylodynamic models, starting with a single ancestor cell. Branches represent cells, internal nodes represent cell divisions, tips represent the sampled cells, branch lengths correspond to time. Second, we simulated CRISPR lineage recordings along the trees. Non-sequential recordings consist of independent targets, where each target acquires an edit at random. In contrast, sequential recordings comprise arrays of target sites, where each array acquires edits in sequential order. We collected barcodes (accumulated edits on target sites) for all sampled cells. Colors indicate different editing outcomes. Third, on each set of barcodes, we applied Bayesian phylogenetic and phylodynamic inference to jointly reconstruct the time-scaled phylogenetic tree and estimate the parameters of cell population dynamics. Fourth, we evaluated the inference by comparing the inferred tree and parameters to the ground truth.

Second, we simulated sequential and non-sequential CRISPR lineage recordings along the phylogenies. We assumed that all target sites are unedited in the ancestor cell and evolve over time. Thus, for each tree and recorder, we obtained an alignment of barcodes. This simulation was done on fixed trees, implicitly assuming that the cell division, death, and differentiation process generating the trees is independent of the barcode evolution process. Indeed, experimentalists aim to insert barcodes which do not alter the cellular dynamics.

Third, from each simulated alignment, we jointly reconstructed the time-scaled cell phylogeny and estimated the parameters of population dynamics in a Bayesian Markov chain Monte Carlo (MCMC) framework in BEAST 2 [31]. Specifically, we applied the editing models *TiDeTree* [17] and *SciPhy* [19] to our simulated non-sequential and sequential CRISPR lineage recordings, respectively, and fitted birth-death sampling models [32–37] to the data.

Fourth, we evaluated phylogeny reconstruction by computing the weighted Robinson-Foulds (wRF) distance [38, 39] between each inferred tree and true tree. Importantly, this metric considers branch lengths in addition to the tree topology and thus evaluates time-scaled cell phylogenies. Additionally, we inspected the posterior distributions of the tree and phylodynamic parameters. We assessed the coverage by computing the fraction of 95% highest posterior density (HPD) intervals which contained the true parameters. Also, we assessed the information gain (when using sequence data in addition to the prior assumptions) by computing the ‘HPD proportion’, i.e., the width of the estimated HPD intervals with respect to prior distributions. Further, we determined the relative bias of the inferred parameters by comparing posterior medians to the true values, and quantified the certainty in the estimates by computing relative HPD widths.

Overall, our workflow served to analyze systematically the phylogenetic and phylodynamic signal in CRISPR lineage tracing data.

### 2.2 Assessing the phylogenetic and phylodynamic signal in CRISPR lineage recordings - the baseline

Initially, we generated cell phylogenies where all cells undergo the same dynamics, and simulated CRISPR lineage recordings under a representative experimental design (baseline).

Specifically, we simulated cell populations growing either by synchronous and regular cell divisions, or in a stochastic manner. In the former, we assumed that the time to division is equal for each cell and defined four models as follows: (1) cells divide and are sampled completely, (2) cells divide and are sampled incompletely, (3) cells divide, die with a fixed probability and are sampled completely, and (4) cells divide, die with a fixed probability and are sampled incompletely (see details in Section 4.1). For stochastic dynamics, we used the constant rate birth-death process [32, 33], where the time to birth (in this context, cell division) and death is exponentially distributed. In total, we obtained 100 phylogenies - 20 per model - with the number of tips on the order of 100. Note that we run additional simulations on tenfold larger trees to assess inference at increasing sample size (for details, see Section S5 in Supplementary Material).

Along the phylogenies, we simulated non-sequential recordings on 20 targets and sequential recordings on 20 tapes of length 5. We used an editing rate of 0.05 so that most, but not all, target sites per cell mutated. Note that even though we applied the same editing rate here, the dynamics of editing between the two systems differ. In a non-sequential recording, all sites are amenable to editing from the start of the experiment, so the effective rate of editing across the whole barcode decreases over time as more sites become edited. In contrast, in a sequential recording, sites within a tape become accessible in order, meaning that early sites can be edited earlier, while later sites become available only after prior insertions. In total, we performed 100 simulations (one per tree) per lineage recorder.

In the inference, we have to specify a barcode evolution model and a tree generation model. For the barcode evolution model, we employed the frameworks *TiDeTree* and *SciPhy* to the non-sequential and sequential barcode evolution, respectively; we used these frameworks for both simulation and inference. The key difference between *TiDeTree* and *SciPhy* lies in their assumptions about barcode evolution, allowing us to directly assess the impact of sequential vs. non-sequential editing. For the cell population growth, i.e. the tree generation, we assumed the commonly employed stochastic birth-death process.

Note that this inference model matches the simulation model perfectly when analyzing birth-death trees, and we expect good estimates of the model parameters given enough data and appropriate priors. However, when analyzing the synchronous cell division trees, there is a model misspecification which may lead to a reduced quality of the estimates. As there is currently no inference model corresponding to the simulation model of synchronous cell divisions, we explored to what extent the cell division and death rates inferred under the birth-death model can be interpreted. In this section, we characterize how well the model parameters can be inferred across dynamics, and in later sections we explore how these estimates change when changing aspects of the simulation.

As shown in Fig. 2, the true editing rate and phylogenetic tree parameters (height and length) had a coverage greater than 80% across all simulations — defined as the frequency with which the true simulation parameter value is contained within the estimated 95% HPD interval — and exhibited only small relative bias. The true cell division and death rates were recovered for all simulations in which the birth-death process generated the phylogenetic tree. However, their recovery was substantially lower for simulations on phylogenetic trees which grew by synchronous cell divisions, as expected due to the model misspecification. In particular, death rates were consistently underestimated for simulations on synchronous trees with cell death, and overestimated for simulations on synchronous trees without cell death. The latter could be explained by the nature of the MCMC chain and the choice of prior distribution on the death rate. We used an exponential distribution, defined on [0, ∞) and continuous, thus, the probability for the chain to take the single value 0 was zero, and the samples were taken from small but positive real numbers. Importantly, all 95% HPD intervals of the posterior estimates approached the true value 0. Consequently, the net growth rate was consistently underestimated, albeit slightly, in the case of synchronous cell divisions without cell death.

**Fig. 2:**
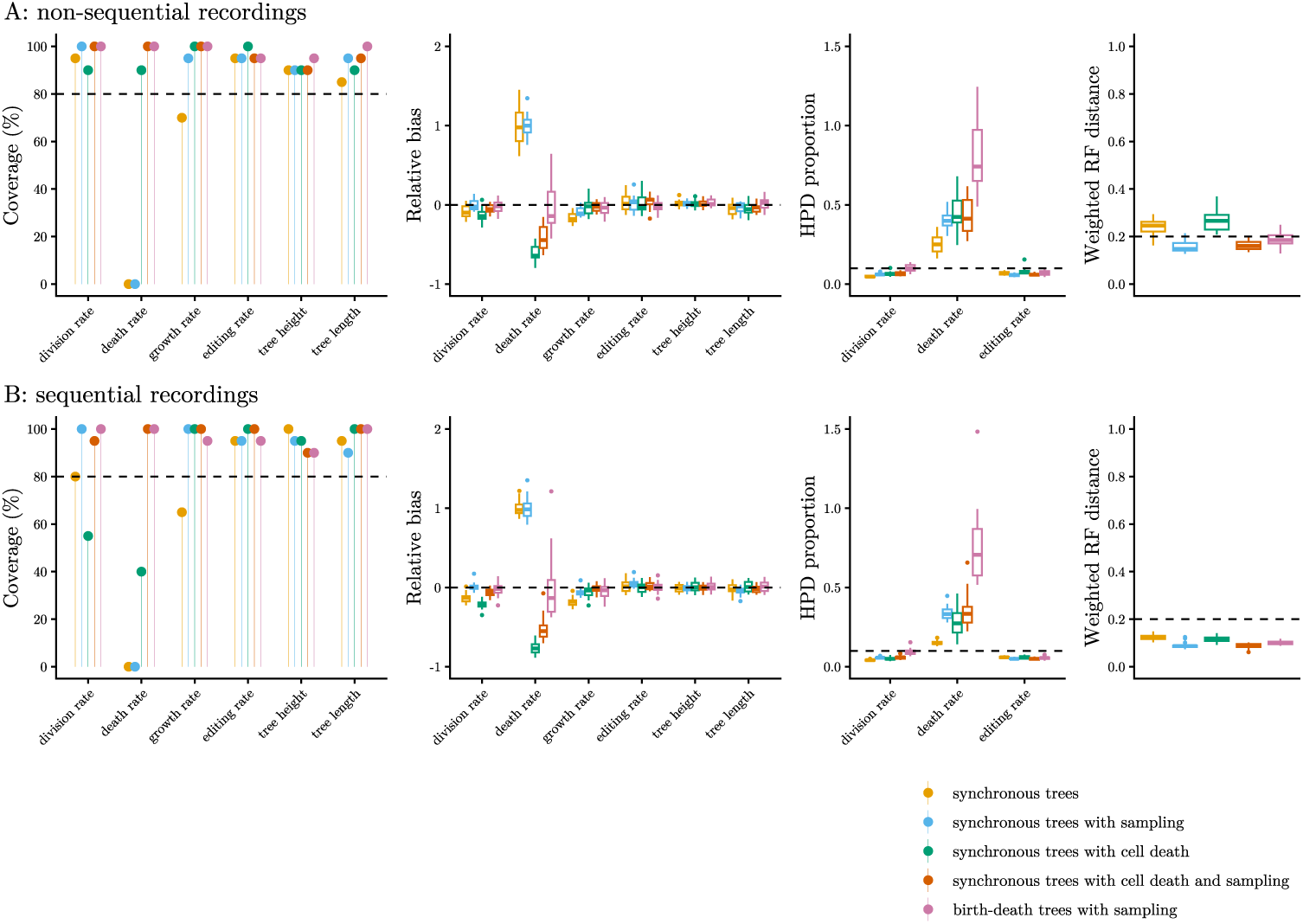
Inference performance on baseline (A) non-sequential and (B) sequential CRISPR lineage recordings. In the left plots, each point indicates the percentage of simulations in which the true parameter was recovered. Next, the box plots show from left to right: the relative bias and HPD proportion of the numerical parameters, and the weighted Robinson-Foulds (RF) distance between the inferred and true trees. The inference results are summarized across 20 simulations per tree generating process (indicated by colors). Dashed lines display thresholds for each metric to facilitate visual comparability.

On average, the posterior distributions of the cell division and editing rates shrank by more than 90% relative to prior distributions, suggesting high confidence in the estimates and strong signal in the data. Compared to these parameters, the uncertainty about the inferred death rates was high (cf. HPD proportion in Fig. 2, and Fig. S1). Further, the wRF distance between the inferred trees and true trees oscillated around 0.2 for non-sequential recordings. The distance was normalized by tree lengths, so it falls between 0 and 1, where 0 indicates identical trees. Hence, the relatively low value of 0.2 indicates that the trees were mostly well resolved, with disagreements involving few splits or short branches which contribute little weight. Notably, the distance was halved (median 0.1) for sequential recordings, which could be due to the overall larger number of target sites in the sequential recorder or due to the sequential edits themselves.

Interestingly, topological reconstruction achieved the highest accuracy for the most regular trees (Fig. S3), that is, the complete, balanced trees produced by synchronous divisions. However, the estimated branch lengths in those trees were overdispersed compared to the true distribution (Fig. S4). This indicates that the birth-death model was not able to fully capture the synchronicity in the timing of cell divisions. Nevertheless, the inferred branch length distributions matched the truth better for sequential recordings, indicating that more informative data yielded estimates of cell division timings that were more robust to model misspecification in the tree prior.

Overall, these results indicate that our simulated CRISPR lineage recordings contained information on both cell relationships and the timing of cell divisions. However, they bore a weaker signal on cell death rates. While cell division estimates were slightly biased for only half the classes of synchronous trees (with complete sampling), the estimation of death rates was more sensitive to model misspecification and biases persisted in all classes of synchronous trees. The observed biases became particularly evident at increased sample size (Figs. S10 and S11). Notably, inference from both larger and sparser samples yielded qualitatively similar results across tree generating processes for both types of CRISPR lineage recorders.

### 2.3 Varying experimental parameters

To evaluate how different experimental parameters affect the inference from CRISPR lineage recordings, we varied the editing rate, the number of editable target sites, and the editing window in our simulations. To ensure that any differences from the baseline are due to the varied parameters, we re-used the 100 phylogenetic trees from the baseline. Thus, we simulated 100 alignments per setting. Fig. 3 illustrates the general trends, whereas Figs. S6 and S7 show differences between the tree generating processes.

**Fig. 3:**
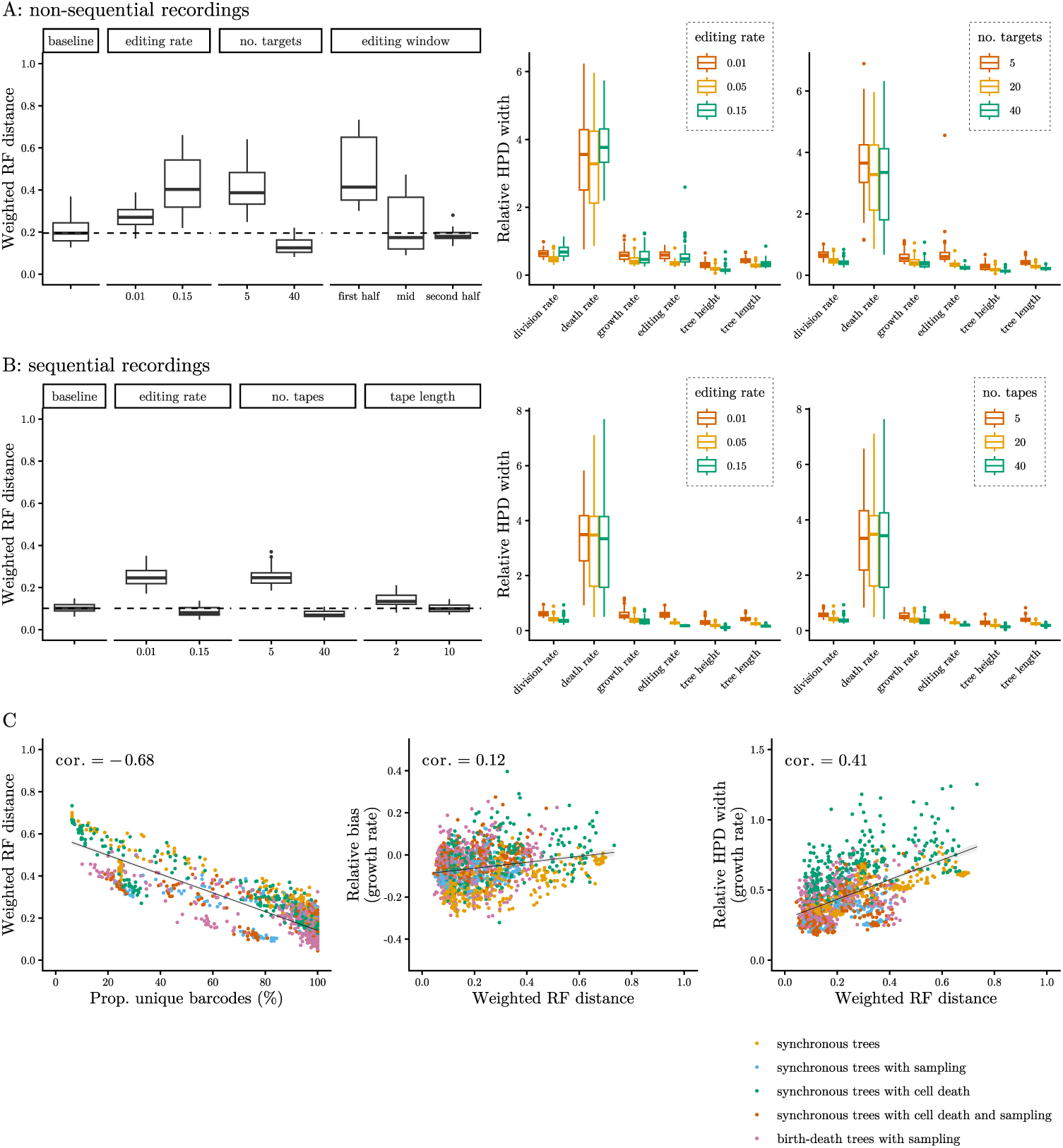
Inference performance on (A) non-sequential and (B) sequential CRISPR lineage recordings with varying experimental parameters. At baseline, the editing rate was 0.05, the recorders carried 20 targets or 20 tapes of length 5, respectively, and editing occurred throughout the entire experiment. Left graphs show the weighted RF distance between the inferred and true trees. Facets indicate the varied experimental parameters, dashed lines display the baseline medians. Right graphs show the relative HPD width for numerical parameters, colored by the editing rate and the number of targets/ tapes used for simulation. The inference results are summarized across 100 simulations per setting. Panel (C) shows associations between barcode diversity (measured by the proportion of unique barcodes per recording), accuracy of phylogenetic reconstruction (measured by the weighted RF distance between the inferred and true trees), and performance of phylodynamic inference (evaluated with the relative bias and HPD width of the growth rate estimates) across experimental scenarios and tree generating processes (indicated by colors). Black lines indicate the linear trends (least squares fit). cor.: Kendall’s *τ* correlation c^4^o^9^efficient.

Of particular interest in CRISPR lineage tracing experiments is calibrating the rate at which target sites accumulate edits. The recorders are characterized by a limited number of target sites, so tuning the editing rate is necessary to obtain sufficiently diverse barcodes at the end of an experiment. A too low editing rate may provide only scarce information, while a too high editing rate leads to recorder saturation before the end of the experiment and missing information on later cell development. Here, we investigated what consequences these scenarios have for phylogenetic and phylodynamic inference.

First, we reduced the editing rate to 0.01 so that less than half of the available targets or tapes per cell mutated. Then, we increased the editing rate to 0.15 so that all targets in the non-sequential recorder and at least the first position in all tapes in the sequential recorder mutated. We observed that phylogeny reconstruction from non-sequential recordings was less accurate for the lower and higher editing rate compared to the baseline. In fact, at the highest editing rate, groups of cells acquired identical barcodes, indicating recorder saturation. Under these conditions, the distance from the inferred to true phylogenetic trees almost doubled. In contrast, reconstruction of the cell phylogeny from sequential recordings was most accurate for the highest editing rate. As target sites in the tapes were activated in order, the tapes did not saturate too early and accumulated information until the end of the experiment.

Next, we decreased and increased the number of targets in the non-sequential recorder, and the number and length of tapes in the sequential recorder. We observed that the inference improved consistently when more target sites were available, as expected. In particular, the average distance between the inferred and true phylogenetic trees has halved for recorders with 20 compared to only 5 targets or tapes, and decreased further for 40 targets or tapes. This result nicely agrees with general statistical theory where adding four times as many observations in the sample reduces the standard error of the mean by a factor of two.

Finally, many systems of interest develop over long time spans over which current technologies cannot sustain editing entirely. Thus, we examined the effect of a limited editing period and its placement during the experiment on parameter estimates. In particular, we restricted the editing window in the less capacious, non-sequential recorder to half of the experimental period. Editing occurred either in the first half, in an interval around the middle, or in the second half of the experiment. In the last case, cell barcodes evolved to the most diverse set. Surprisingly, phylogeny reconstruction from barcodes edited in the middle or second half of the experiment was on average as accurate as reconstruction from barcodes edited throughout the entire experiment, as measured by the weighted RF distance. This result might be affected by the timing of the cell division events. As more cell divisions occur towards the end of the experiment, the number of splits increases, and later editing appears better under that metric. Generalized RF metrics, measuring tree topology only, indicate that phylogeny reconstruction was less accurate when editing was restricted to the first or second half, compared to the baseline, but was similarly accurate when editing occurred in an interval around the middle of the experiment (cf. Fig. S7).

Again, we evaluated the inference of cell division and death rates alongside cell phylogenies. Notably, its performance differed much more subtly across the experimental parameters. In general, we observed that HPD intervals of the cell division and death rates became narrower as the capacity of the recorders increased, indicating higher confidence in the estimates. Also, relative HPD widths mirrored the trends in phylogeny reconstruction for the different editing rates, and numbers of targets or tapes (as illustrated in Fig. 3 right and Fig. S6).

Across scenarios, we observed a negative correlation between barcode diversity per recording and the weighted RF distance between the inferred and true trees (Kendall’s *τ* = −0.68, 3C). That is, the more cells had a unique pattern of accumulated edits across target sites, the more accurate was the tree inference. More precise and reliable branching times, in turn, propagated to the phylodynamic estimates (cf. Fig. S5). Accordingly, the weighted RF distance correlated moderately with the relative HPD width of the inferred growth rate (*τ* = 0.41), but only weakly with its bias (*τ* = 0.12). Crucially, once nearly all cells acquired unique barcodes, the lineage relationships could be resolved almost perfectly. Further accumulation of edits resulted in only minor improvements in phylogenetic and phylodynamic inference. For example, the sequential recorder reached 40% saturation at an editing rate of 0.05 (on average, 40 out of 100 sites were edited per recording) and 90% saturation at an editing rate of 0.15, more than doubling the number of informative sites (cf. Table S2). In both cases, the resulting barcodes were highly diverse and the inferred phylogeny was very accurate (average weighted RF distance of 0.1 and 0.09, respectively, cf. Fig. S7). However, biases in the inferred branch lengths and population-dynamic estimates due to phylodynamic model misspecification persisted, even as information content increased (cf. relative biases of division and death rates in Fig. S6, KS distances in Fig. S7, and Fig. S8).

In summary, recorders with more editable target sites and an editing rate tuned to recorder capacity – generating more diverse barcodes – carried a stronger phylogenetic and phylodynamic signal. It is also notable that some parameters, such as the cell division rate, can be robustly estimated even when the editing window is halved.

### 2.4 Evaluating sequential editing

While the non-sequential recorder is limited to one edit per target, the sequential recorder has multiple editable positions per tape. At baseline, we saw that the sequential recorder accumulated more edits throughout the experiment, so more diverse barcodes arose. Therefore, phylogeny reconstruction improved and uncertainty in the parameter estimates decreased. But did the inference improve only due to an increase in recorder capacity, or due to the sequential acquisition of edits per se?

To test this, we simulated a scenario where we held everything essentially equivalent and contrasted 20 non-sequential sites with 20 sequential sites (one tape of length 20). For the non-sequential recorder, we could reuse the baseline. For the sequential recorder, we used the same number and distribution of possible editing outcomes as for non-sequential recordings. We further tuned the editing rate to 0.45, such that the average number of edits per barcode and the diversity of barcodes were comparable to the non-sequential recordings. Note that the editing rate *r* is defined per active target site. In the non-sequential model, all *m* target sites are active from the start on, while in the sequential model, target sites in a tape are activated one by one. Hence, the overall editing rate in the non-sequential model is initially *r* · *m* and decreases over time until all targets are edited. In contrast, in the sequential model, it is initially *r*, constant over time, and changes to 0 only when all target sites are edited. Therefore, tuning the editing rate was necessary to reach a similar amount of edits per cell in the two types of recordings.

Importantly, phylogeny reconstruction, as measured by weighted RF distance, improved significantly when edits accumulated in order (one-sided, paired Wilcoxon signed rank test, *V* = 4754*, p <* 0.001, see Fig. 4A). However, the inference of phylodynamic parameters did not change substantially (Fig. 4B). Evaluating branch lengths separately from tree topology revealed that sequential editing primarily improved the reconstruction of lineage relationships, while the temporal aspect of the trees remained largely unaffected. Consequently, sequential editing did not consistently increase the accuracy nor reduce the uncertainty in the estimated cell division and death rates (see Fig. S9 for results across tree generating processes).

**Fig. 4:**
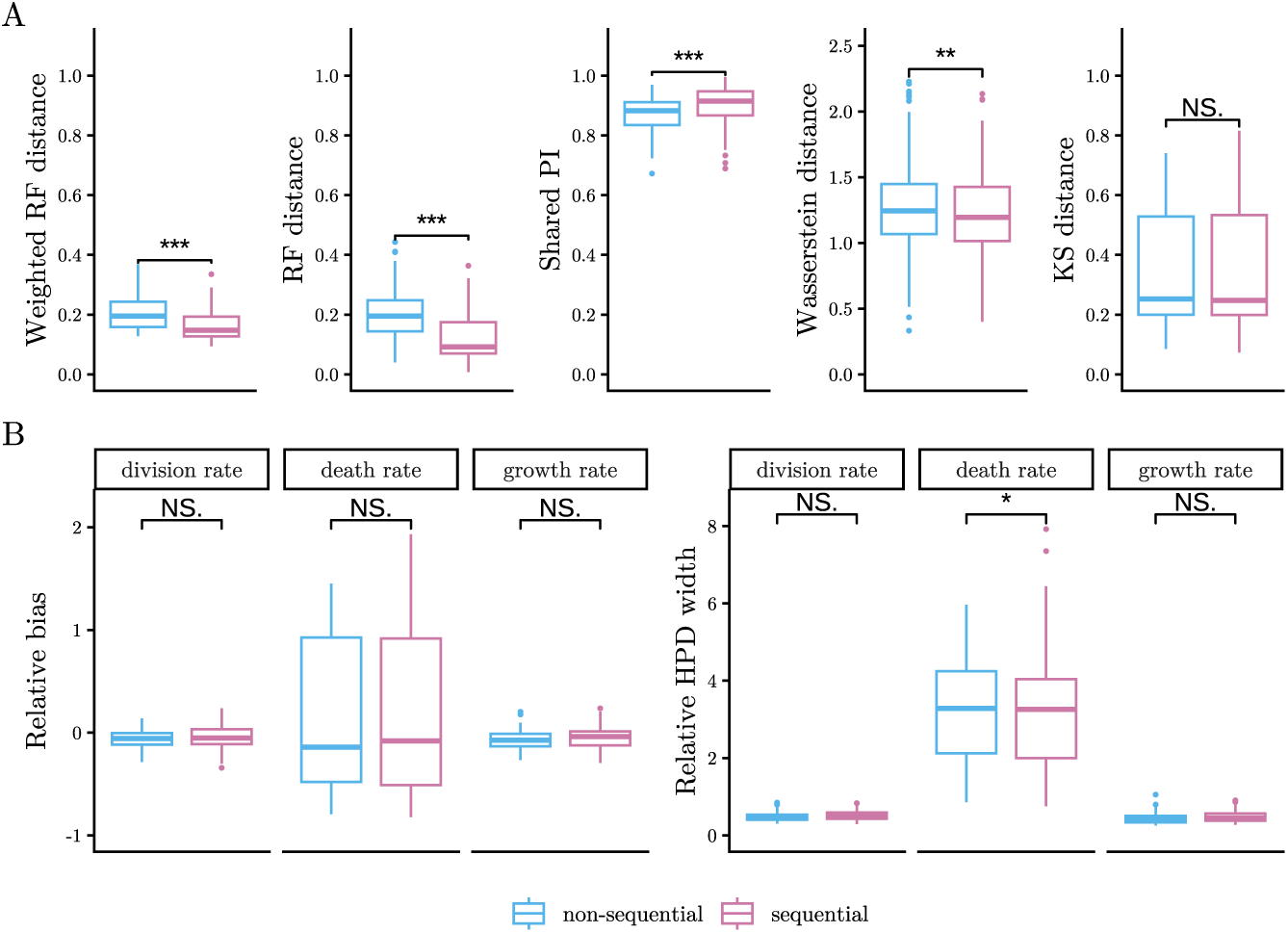
Comparison of the sequential and non-sequential accumulation of edits in CRISPR recorders. Panel (A) compares the inferred and true trees using weighted RF distance, topological RF distance, Shared Phylogenetic Information (PI), as well as Wasserstein and Kolmogorov–Smirnov (KS) distance between branch length distributions. Panel (B) shows the relative bias and HPD proportion of the inferred cell division, death, and growth rates. The inference results are summarized across 100 simulations per setting (including both simulated synchronous and birth-death trees). Statistical significance was assessed by one-sided, paired Wilcoxon signed rank test, ***: p-value *<* 0.001, **: p-value *<* 0.01, *: p-value *<* 0.05, NS.: not significant.

### 2.5 Filtering out noisy data

In practice, CRISPR lineage recordings contain substantial portions of noisy or missing barcodes, arising mainly from two mechanisms [40]. First, the CRISPR editing process can lead to target silencing, resulting in heritable loss of lineage information. Second, CRISPR barcodes are prone to technical dropout during single-cell sequencing. During analysis, a common strategy is to filter out target sites and cells with high rates of missing or unreliable barcodes, and then reconstruct lineage trees for the remaining cells. This filtering procedure differs fundamentally from random sampling, which assumes that cells are independently and uniformly drawn from the population. Especially in the case of target silencing, entire clades of cells may be excluded from analysis, potentially biasing the results in a systematic manner.

To assess the impact of missing data on the inference, we simulated CRISPR lineage recordings along larger trees at baseline experimental parameters, incorporating both target silencing and sequencing dropout. This resulted in 40 − 60% of missing barcodes per recording. We then filtered the data, as currently done in practice [11], by retaining the largest subset of targets (or tapes) for which at least 10% of cells contained complete barcodes. Effectively, we could then infer trees and phylodynamic parameters for subsets of cells (of size 35 − 250) with 5 out of 20 targets (or tapes) on average retained per recording. Crucially, the inferential models did not explicitly account for the error mechanisms and the filtering procedure.

Nevertheless, the true parameters – cell division and death rates, editing rate, tree heights and lengths – were recovered in at least 90% of simulations (only death rates of 0 could not be captured, as before). Interestingly, filtering affected the inference of cell division rates more strongly in the case of synchronous than birth-death trees, leading to slight systematic overestimation of this parameter (Fig. 5).

**Fig. 5:**
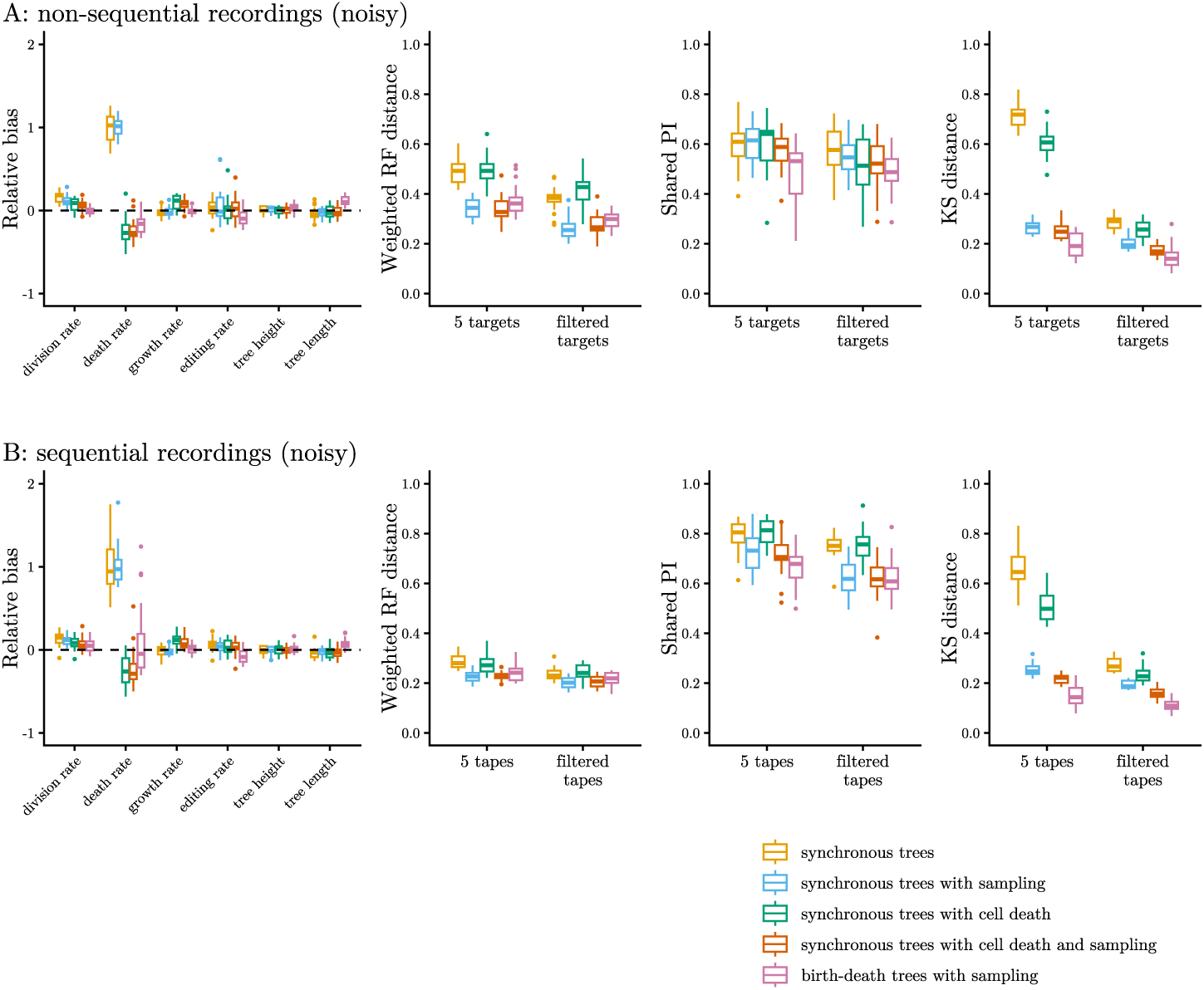
Inference from noisy (A) non-sequential and (B) sequential CRISPR lineage recordings. The left plots show the relative bias of parameters inferred from filtered barcodes. Next, the panels compare tree inference from filtered barcodes (consisting on average of 5 targets/ tapes per simulation) to recordings with 5 editable targets/ tapes (without errors). The plots show dis(similarity) metrics between the inferred and true trees, from left to right: weighted RF distance, Shared Phylogenetic Information, and Kolmogorov–Smirnov distance between branch length distributions. The results are summarized across 20 simulations per tree generating process (indicated by colors).

Further, we compared tree inference from filtered barcodes to recordings, in which only five targets or tapes were available for editing and no errors occurred. On average, topologies reconstructed from filtered barcodes were less accurate (median Shared Phylogenetic Information of 0.53 vs. 0.59 for non-sequential, and 0.67 vs. 0.74 for sequential recordings, cf. Fig. 5, third panel). However, under the weighted RF distance – which also accounts for differences in branch lengths – the inferred trees did not deviate more strongly from the true trees. In particular, when a filtering step preceded inference, the branch length distributions of the inferred and true synchronous trees were more similar, as quantified by the Kolmogorov–Smirnov distance. The primary reason is that filtering effectively induced non-uniform sampling of lineages, making branches in the synchronous trees more irregular and more compatible with the birth-death model used for inference (cf. Fig. S13).

Overall, population-dynamic parameters were inferred with comparable accuracy from filtered recordings, despite retaining only ≈ 2.5% of the data, albeit at the cost of reconstructing lineage trees for only a small subset of cells.

### 2.6 Inferring cell differentiation dynamics

A key interest of developmental biologists are the differentiation dynamics of different cell types. To investigate the inference of these dynamics, we simulated non-sequential and sequential CRISPR lineage recordings with baseline experimental parameters along multi-type birth-death trees [34, 35]. We considered two prototypical dynamics of cell type transitions, terminal and chain-like, of three different cell types (1, 2 and 3) from a single progenitor type 0 (see details in 4.1). From the recordings with end-point cell type annotation, we jointly inferred time-scaled cell phylogenies, cell type-specific division and death rates, as well as cell type transition rates (Fig. 6). We then applied a stochastic mapping algorithm, as implemented in the BEAST 2 package *BDMM-Prime* [37], to reconstruct ancestral type changes along the lineage trees (see Fig. S14 for example multi-type trees and their inferred counterparts).

**Fig. 6:**
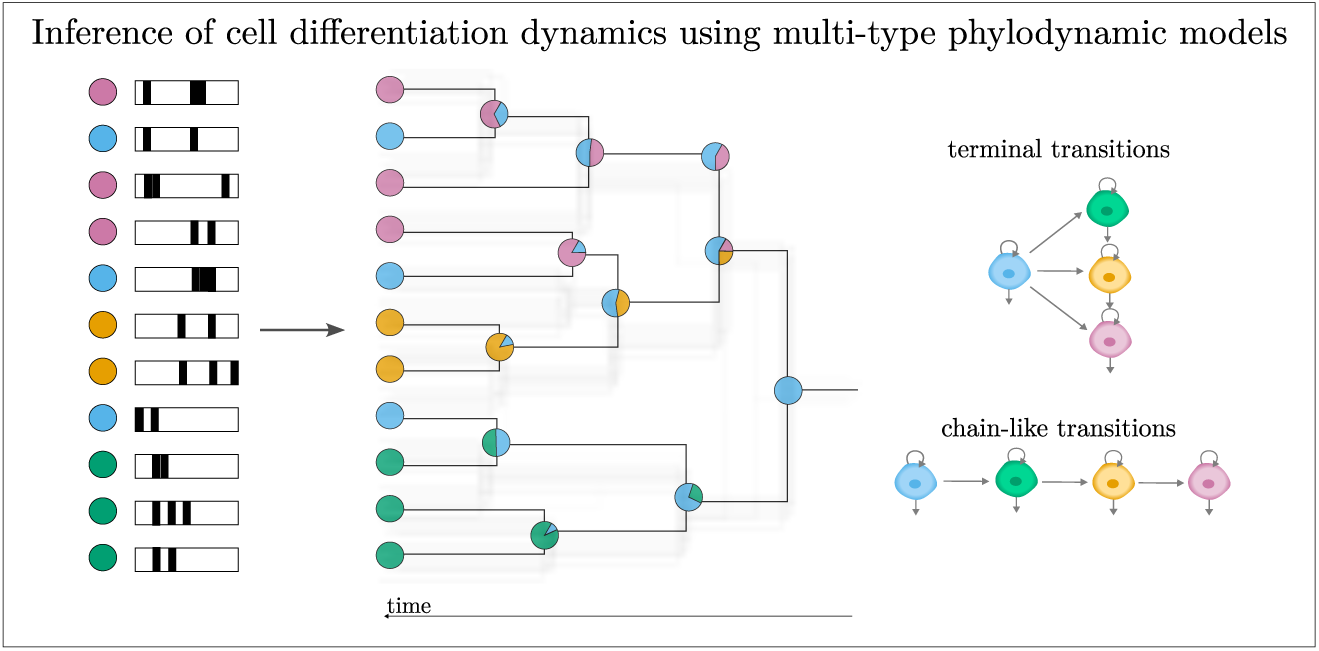
Schematic of the multi-type inference. From simulated CRISPR lineage recordings with end-point cell type annotation, we inferred time-scaled phylogenetic trees with ancestral cell type probabilities. Additionally, we estimated cell-type specific division and death rates and cell type transition rates, assuming two prototypical cell type transition dynamics.

Importantly, the coverage of all parameters was above 80% for both recorders and both cell type transition dynamics (Fig. S17). In particular, the inferred transition rates varied around the true values for most simulations, and their HPD intervals were much narrower than the prior distribution (median HPD proportion 0.35). For trees with terminal transitions, uncertainty was lowest for transition rate to type 1 and highest for transition rate to type 3. Given that we simulated the trees with a transition rate to type 1 two times higher than the transition rate to type 3, we hypothesized that the higher the true transition rate, the more cells of the corresponding terminal type were sampled, and the higher was the signal for inference.

For both recorders, the transition rate from type 1 to type 2 was overestimated almost tenfold for some trees with chain-like transitions (see Fig. 7 middle row). A closer look at these trees revealed that none of them contained a single tip of type 1 (Fig. S15 right). This implies that transitions from type 0 to type 1, or transitions from type 1 to type 2, or both, had occurred very fast such that no cells of type 1 were sampled at the end of the experiment. In the inferred trees, transitions to type 2 were placed closer to the process origin (Fig. S15 left). This temporal shift is consistent with the inflated rate estimates, as the expected waiting time for a transition under the multi-type birth-death model is inversely proportional to its rate. Earlier transitions, in turn, are more likely to produce clades dominated by a single cell type (in this case, type 2). Note, however, that the inflated rates exhibit very wide 95% HPD intervals, indicating high uncertainty in the estimates.

**Fig. 7:**
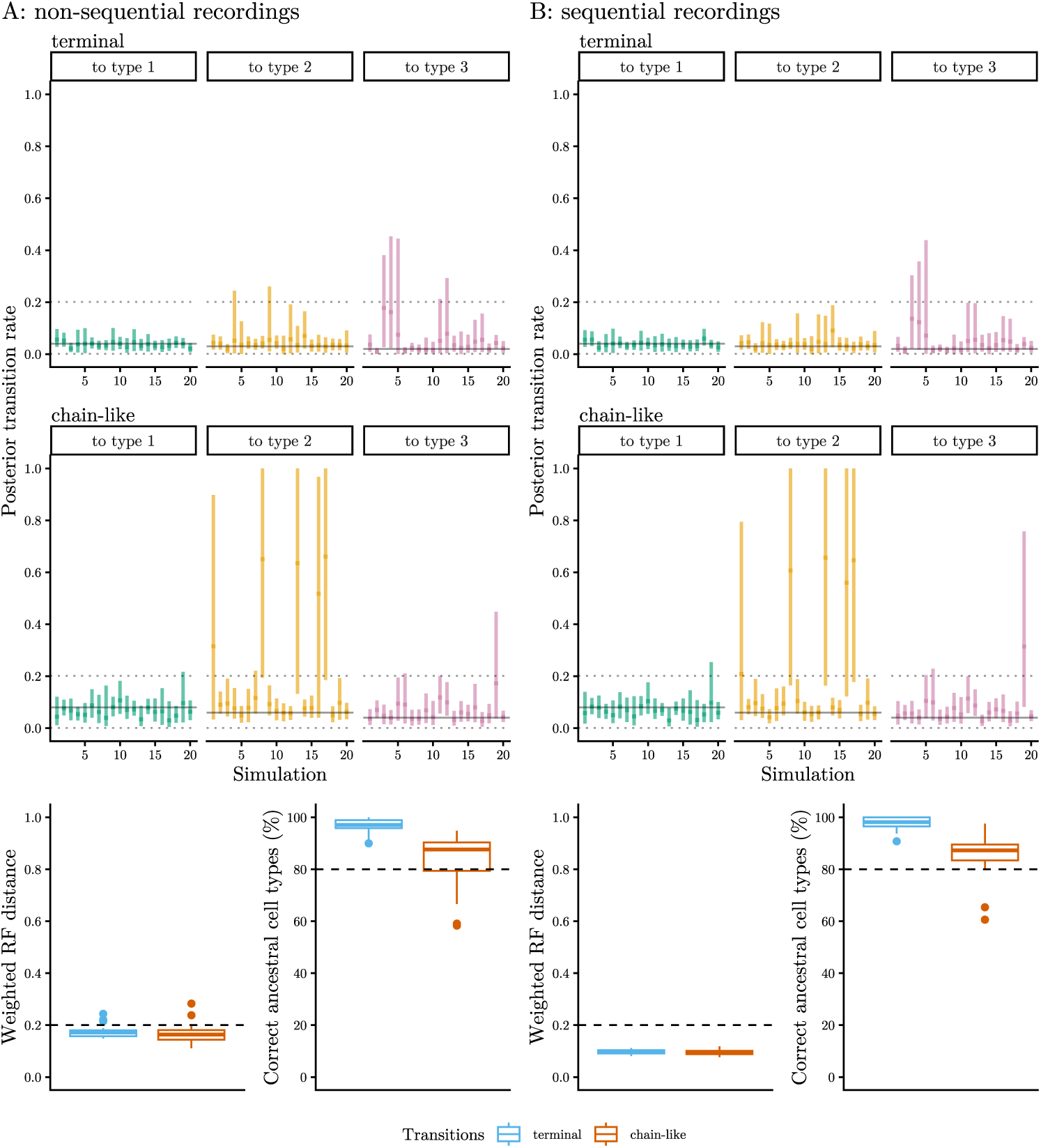
Inference from non-sequential (A) and sequential (B) recordings on multi-type birth-death trees. The top two panels show the medians (dots) and 95% HPD intervals (lines) of the inferred transition rates per simulation. Solid horizontal lines indicate the true values, dashed horizontal lines indicate the 95% credible interval of the prior distribution. The bottom panels show the weighted RF distances between the inferred and true trees, and the percentage of correctly estimated ancestral cell types for the two transition dynamics. The metrics are summarized across 20 simulations per recorder and cell type transition dynamics. Dashed lines display thresholds for comparability.

Notably, the topology and branch lengths of multi-type phylogenies were inferred comparably to those of single-type phylogenies (Fig. 7 last row). Additionally, cell types were correctly inferred for more than 80% of ancestral cells across settings. They were consistently inferred better in trees with terminal transitions (median 97.8%) than those with chain-like transitions (median 87.4%). This could easily be explained by the observation that each cell could be either of the same type as its descendant or of type 0, whereas more possibilities exist in the case of chain-like transitions. Additionally, statistics for each cell type – the number of transitions during the tree generating process, the timing of first transition, and the total time spent in each type (i.e., the sum of branch lengths per type) – were captured well across simulations (Fig. S16).

Taken together, the rates and trajectories of cellular differentiation were inferred robustly from CRISPR lineage recordings alongside cell relationships when enough cells of the different types were sampled.

## 3 Discussion

We have carried out a simulation study to assess the inference of single-cell phylogenies and population dynamics from CRISPR lineage tracing data. In line with Salvador- Martínez et al. [14] and recent theoretical analyses [15, 16], we find that phylogeny reconstruction largely depends on the capacity of CRISPR lineage recorders. Principally, recorders with more target sites can accumulate more edits throughout the experiment and produce more informative barcodes. Crucially, when the editing rate is too low or too high for a given number of target sites, not all cell divisions can be recorded which leads to only partially resolved cell phylogenies. Our results indicate that less accurate phylogeny reconstruction goes along with less certain estimation of phylodynamic parameters.

The study was designed to compare non-sequential and sequential CRISPR lineage recorders. As expected, replacing targets with tandem arrays of target sites increases recorder capacity, and thus improves the inference. In our simulations, we have accounted for the fact that CRISPR–Cas9 machinery in non-sequential recorders produces diverse, largely random scars, whereas prime editors in sequential recorders introduce fewer but predefined insertions. Nevertheless, when on average two to three positions in the arrays had accumulated insertions, barcode diversity at the final time point exceeded that observed when each target contained only a single scar. Quantitatively, this corresponded to an approximately twofold reduction in the weighted Robinson–Foulds distance between inferred and true trees. Importantly, assuming comparable editing outcomes, we find that the order of edits in sequential recordings carries additional information that significantly improves phylogeny reconstruction in terms of topology. However, sequential editing does not substantially improve the temporal resolution of phylogenies, and therefore provides little additional benefit for the inference of phylodynamic parameters.

Further, we have observed that non-sequential recorders saturate faster than sequential recorders at the same editing rate per active target site. This is because, in the non-sequential setting, all target sites are simultaneously active from the start of the experiment, resulting in a higher initial editing activity across the entire barcode and hence faster saturation. In contrast, sequential recorders activate sites gradually, limiting the number of editable sites at any given time. As a result, sequential recorders accumulate edits more slowly and can track lineages over longer periods. One strategy to prevent saturation of a non-sequential recorder during the experiment, is restricting the editing window to the most interesting period of development. Previously, it has been reported [4] that early editing facilitates the recognition of specific clones in a cell population, but inferring relationships among cells within the clones is difficult. Consistent with this, we find that editing in only the first half of the experiment leads to less well resolved phylogenetic trees. Our results indicate, however, that phylogeny reconstruction can be almost as accurate when editing lasts half the time in an interval around the middle or the second half of the experiment as when it spans throughout the entire experiment. However, this observation likely depends on our simulation setup, where few cell divisions occur early on, and may not generalize to scenarios where most divisions happen at the beginning of the experiment.

In our simulations, we have additionally explored the effects of filtering out noisy data prior to inference – a step particularly relevant to real-world CRISPR lineage recordings, since ambiguous or missing barcodes are abundant in empirical datasets. Such filtering effectively reduces the number of target sites that can be reliably aligned across cells and limits phylogenetic reconstruction to only a subset of the sampled population. Explicitly incorporating error processes into the mechanistic, inferential models of CRISPR editing (as demonstrated in [40]) would circumvent the need for extensive filtering, thereby preserving a larger proportion of the dataset for analysis, reducing the sampling bias, and leveraging the additional phylogenetic signal which arises from heritable target silencing.

Most importantly, our study has revealed how well the parameters of population dynamics can be estimated from CRISPR lineage recordings using the Bayesian inference framework. Notably, we used different cell population models (either models based on synchronous cell divisions or a birth-death process) for simulation and inferred relatively accurate cell division rates across dynamics despite model misspecification. Thus, we find that the recordings carry a strong signal on the rates of cell division, as evidenced by high accuracy and certainty of the estimates across scenarios.

However, our results indicate that signal on the rates of cell death is weak and their estimation is sensitive to model misspecification. Theoretical considerations on the inference of birth and death rates from molecular phylogenies can explain this result. It has been shown that death events leave a characteristic signature in the shape of phylogenetic trees, and can be derived from the increase in the number of lineages through time [41, 42]. However, in small populations, the number of lineages through time is very noisy, and often, not sufficiently many death events occur to reliably estimate the death rate [43]. Furthermore, the quantification of death rate becomes more challenging when only a sample of cells is analysed [41]. Thus, while the signal for death rates is weaker than for cell division rates, this is in line with the use of phylodynamic methods in adjacent fields and not a characteristic of lineage tracing data per se.

Another interesting result is that cell type-specific division rates and differentiation rates can be estimated from CRISPR lineage recordings, when enough cells of each type are sampled (and given no model misspecification). In our simulations, at least one cell of each type had to be present in the dataset to inform the type-specific phylodynamic parameters, and we required cells to be sampled with the same probability (irrespectively of the type) at the end of the experiment. This finding has implications for developing appropriate procedures to sample cells in real-life experiments. Sampling schemes on experimental replicates should capture sufficiently many cells of all types. It should be explored in further simulations how non-uniform sampling and sampling biases of certain cell types affect the inference. This might be particularly relevant when the real cell type transitions are much more complex than assumed in our simulations, and when more different cell types arise during development, some of which abundantly and others rarely.

In wider perspective, applying multi-type phylodynamic models to single-cell data opens the door to characterizing cell differentiation dynamics in real time. Novel lineage tracing technologies (e.g., [7, 11, 44]) are typically combined with gene expression profiling of end-point samples, enabling cell type annotation along lineage reconstruction. This study demonstrates that, in principle, phylodynamic analysis of such data allows for the quantification of cell type transition rates within the population. More importantly, it enables the inference of ancestral cell type transitions and their timing along cell lineages, addressing the limitations of snapshot-based differentiation trajectory analyses that do not take cellular ancestry into account [3].

While our simulations focused on mimicking cellular differentiation in early development, the multi-type approach is equally applicable to characterize cell population dynamics in disease, particularly clonal dynamics in cancer (similarly to [28]). Within the Bayesian framework, structured phylodynamic models have recently been applied to single-cell genomic cancer data to infer malignant population dynamics over time [45]. The multi-type birth-death model employed here generally allows for unordered transitions between states, extending its utility beyond directed differentiation scenarios. Crucially, it can account for type-specific birth and death rates, which has been shown to be important for accurately recovering transition rates between subpopulations [46]. This approach has proven successful in epidemiology and macroevolution – for example, in quantifying viral transmission across geographic compartments [35, 36] – and could be readily adapted to model transitions between plastic cell states or competing tumor clones, and in the long term, enable the inference of transition graphs beyond merely estimating transition rates.

In future research, more realistic models of cell population dynamics should be developed and evaluated. Such models would overcome biases in estimating the parameters of cell population dynamics due to model misspecification. Our analysis has shown, for example, that estimates of the death rate were biased when we simulated synchronous and regular cell divisions, but inferred the rates of cell division and death under the stochastic birth-death model. More realistic models might further consider, for instance, that cell divisions are initially regular and synchronous, and later become asynchronous and stochastic. They should also account for more complex dynamics of cellular differentiation, and varying rates of cell division and death across types, lineages, and over time. Inference models that faithfully describe cell population dynamics would improve phylodynamic inference from empirical lineage tracing data.

In our simulation study, we did not consider several factors that might affect the phylogenetic and phylodynamic inference from CRISPR lineage recordings. For example, in empirical experiments, the rate of editing can change over time [8], as it is challenging to maintain the desired level of activity of the editing enzyme over long periods of time [4]. Furthermore, technical issues other than target silencing and dropout might occur [1, 14], resulting in CRISPR editing model misspecification. Future studies should account for such factors in simulations and evaluate their impact on inference.

Our workflow has faced the computational limitation of high runtime complexity. Each MCMC chain required at least a few hours to collect sufficiently many samples and converge to the posterior distribution. Inference with the multi-type phylodynamic model lasted up to three weeks. Scaling Bayesian inference methods to analyze lineage recordings on populations consisting of thousands or even millions of cells is a core challenge. Recent computational advancements in the field [47, 48] offer promising avenues for handling such large datasets. On the other hand, it remains an open question how many cells are required to get accurate estimates of cell division, differentiation, and death rates, and thus, to what number of cells the methods need to scale. In this study, we analyzed relatively small trees and, despite data sparsity, recovered the rates for most simulations. Theoretically, when the model adequately reflects the true generative process, increasing the number of cells improves the Bayesian inference, in the sense that posterior distributions concentrate near the true parameters and trees. In other words, for larger trees, the rate estimates become more precise. However, if the model fails to capture the true dynamics, more data may lead to over-confident but biased posteriors - again highlighting the need for developing proper models.

We envision that this study provides guidance for practitioners of the CRISPR lineage tracing techniques and helps to inform experimental design aimed at generating biologically meaningful, information-rich data. Our findings indicate that sequential recorders tendentially contain stronger phylogenetic and phylodynamic signal than non-sequential recorders. For both recorder types, we generally recommend performing forward simulations – the fast and computationally inexpensive step in the pipeline – before running experiments. These simulations should account for the growth dynamics of the cell population of interest, the CRISPR editing machinery, the process of sampling cells for sequencing at the end of the experiment, and the filtering step. Simulated recordings can then be used to assess the diversity of barcodes expected in the sampled cell population. Barcode diversity, in turn, serves as a useful proxy for the amount of signal in the data and the expected performance of phylogenetic and phylodynamic inference. This procedure can be used to determine the required number of target sites and calibrate the editing rate to recorder capacity and recording duration for robust inference.

Altogether, phylogenetic and phylodynamic inference from CRISPR lineage recordings is promising, but faces various challenges, as we have shown in our simulations. Beyond extending the mechanistic models of CRISPR editing for reliable phylogenetic inference, establishing phylodynamic models which accurately represent various cell population dynamics, such as synchronous cell divisions, is crucial for overcoming persisting biases due to model misspecification. Moreover, high runtime and applicability to only small samples are currently the major computational bottlenecks of the inference framework which should be addressed as genetic lineage tracing technologies and analysis methods advance. Going forward, it will be exciting to integrate genetic lineage tracing data with other molecular single-cell measurements, with the goal of establishing more comprehensive models of and gaining insights into cellular development.

## 4 Methods

### 4.1 Simulation of phylogenetic trees

We started each tree generating process with a single cell at time *t* = 0 and let the cell populations evolve until time *T* = 40, measured in arbitrary time units. We generated time-scaled phylogenetic trees in which branches represent cells, internal nodes represent cell divisions, tips represent contemporaneous cells and branch lengths correspond to time.

#### Homogeneous cell populations

In the first set of simulations, we assumed all cells share the same population rates of cell division, death and sampling, and generated phylogenetic trees in R using the packages *ape* [49] and *TreeSim* [50]. We considered five population dynamics: First, we assumed that cells divide synchronously in regular time intervals of length 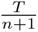, where *n* is the number of cell division time points, and all their descendants survive (*synchronous trees*). Next, we added the possibility that cells die with a fixed probability *p_d_* before the synchronous cell division time points (*synchronous trees with cell death*). For both processes, we then sampled living cells from the population at the end of the process with sampling fraction *ρ <* 1. The resulting *synchronous trees with sampling* or *synchronous trees with cell death and sampling*, respectively, contained only lineages of the sampled cells. Finally, we generated trees using the constant rate birth-death sampling model [32, 33], parameterized by a birth (cell division) rate *β*, a death rate *δ*, and sampling probability *ρ* with which cells at time *T* = 40 are sampled. In these *birth-death trees with sampling*, cells divided or died in a stochastic manner.

#### Heterogeneous cell populations

In the second set of simulations, we generated cell phylogenies under the multi-type birth-death branching model [34] using the package *BDMM-Prime* [37] in *BEAST v2.7.4* [31]. Here, we assumed that cell populations consist of multiple types, each having its own birth (cell division) rate *β_i_* and death rate *δ_i_*. Transition rates *γ_ij_* (in *BDMM-Prime* called migration rates) describe the rate of cells changing from one type *i* to another *j*. We considered four cell types such that *i, j* = 0, 1, 2, 3. Again, we sampled cells at the end of the process, at time *T* = 40, with sampling probability *ρ*, i.e. each cell was equally likely to be sampled.

Based on experimental observations, we considered two prototypical dynamics of cell type transitions. The first one was motivated by a study on *C. elegans* which reported that cell type changes can be abrupt and many distinct terminal cell types arise from progenitor cells [51]. Hence, in trees with *terminal transitions*, cells of type 0 could transition to type 1, 2 or 3, but all descendants of cells of type 1, 2, 3 were of the same type.

The second dynamics was inspired by research on mouse embryonic stem cells which revealed a stochastic, chain-like network of cell-state transitions [52]. Thus, in trees with *chain-like transitions*, cells of type 0 could only transition to type 1, cells of type 1 to type 2, and cells of type 2 to type 3. In this study, we considered only irreversible cell type transitions.

We chose the parameters of all population dynamics described above such that the expected number of sampled cells, corresponding to the number of tree tips, was roughly 100. Although this number is relatively small for single-cell lineage tracing experiments of organism development, we generally expect an increase in signal and improvement in inference for more data (given adequately specified models and fixed sampling proportion [53]). We performed selected simulations and analyses on larger trees to verify that this statement holds for our data (details in Section S5 in Supplementary Material). By operating on small trees, we reduced the computational complexity of the Bayesian inference, and implicitly assessed how much information can be extracted from sparse observations. Our choice of parameters was also guided by simulations in previous studies [17] and an empirical observation of cell development from mouse embryonic stem cells [54]. The parameters of the trees are summarized in Tables 1 and 2 below.

**Table 1:**
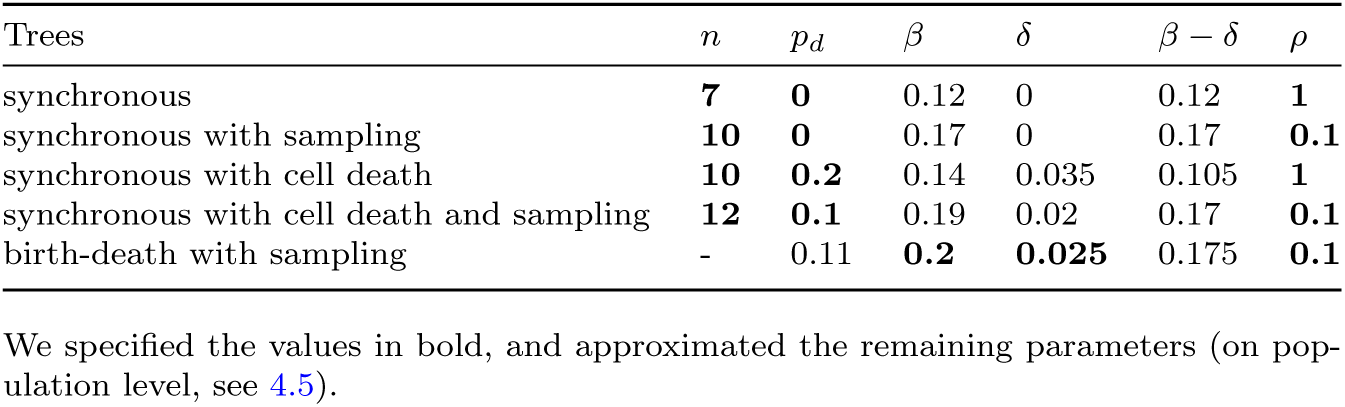
Parameters for simulating homogeneous phylogenetic trees.

**Table 2:** Parameters for simulating heterogeneous phylogenetic trees.

| Parameter | Value |
| --- | --- |
| division rate | $\beta = (\beta_i)_{i=0,1,2,3} = (0.2, 0.18, 0.18, 0.18)$ |
| death rate | $\delta = (\delta_i)_{i=0,1,2,3} = (0.01, 0.03, 0.03, 0.03)$ |
| transition rate | $\gamma_{\text{terminal}} = (\gamma_{ij})_{i,j=0,1,2,3} = \begin{pmatrix} 0 & 0.04 & 0.03 & 0.02 \\ 0 & 0 & 0 & 0 \\ 0 & 0 & 0 & 0 \\ 0 & 0 & 0 & 0 \end{pmatrix}$<br>$\gamma_{\text{chain-like}} = (\gamma_{ij})_{i,j=0,1,2,3} = \begin{pmatrix} 0 & 0.08 & 0 & 0 \\ 0 & 0 & 0.06 & 0 \\ 0 & 0 & 0 & 0.04 \\ 0 & 0 & 0 & 0 \end{pmatrix}$ |
| sampling proportion | $\rho = 0.1$ |

For comparability across all simulations, we selected trees with 20 to 200 tips. In the multi-type case, we assigned slightly different cell division, death and transition rates to the different types. From phylogenies with terminal transitions, we selected trees in which the sampled population consisted of at least one cell of type 1, 2 and 3. From phylogenies with chain-like transitions, we selected trees in which at least one sampled cell reached type 3.

### 4.2 Simulation of barcodes

We simulated the cumulative acquisition of edits in CRISPR lineage recorders in BEAST 2 using the packages *TiDeTree* [17] and *SciPhy* [19].

The frameworks *TiDeTree* and *SciPhy* model the evolution of each target as a continuous time Markov chain on the state space of all possible editing outcomes. The unedited state is encoded as 0, the possible edits in *TiDeTree* (scars) are indexed by 1*, …, S* and the possible edits in *SciPhy* (insertions) are indexed by 1*, …, I*.

#### Non-sequential recordings

*TiDeTree* models the following CRISPR-Cas9 editing process: At time *t* = 0, a single cell exists with *m* unedited targets at independent genomic loci. At time 0 ≤ *t_e_ < T*, where *t_e_* denotes the editing height and *T* the experiment duration, the editing window begins by injecting the editing reagents or inducing their expression. The editing reagents are guided to target sites which, consequently, acquire irreversible indels at a constant editing rate *r*. The scarring probabilities *s*_1_*, …, s_S_*indicate the relative frequency of each edit, also called scar, appearing on a target. In subsequent rounds of cell division, the accumulated edits are passed on to descendant cells. Editing may be suspended after a time interval Δ*t_e_*, the editing duration, for example, when the editing reagents degrade. At the end of the experiment, at time *T*, a subset of cells is selected for single-cell sequencing and the accumulated edits at target sites (barcodes) are read out.

#### Sequential recordings

In contrast, *SciPhy* models ordered editing in a CRISPR lineage recorder with a prime editor. The ancestor cell at time *t* = 0 contains *k* tagged tandem arrays (tapes) of *l* target sites. At start, all but the first targets in each tape are inactive. Once the editing reagents are guided to a target site and an editing event takes place, the position of the active target is shifted by one unit along the array. Editing outcomes are, in this case, irreversible short template-based insertions that occur with insert probabilities *i*_1_*, …, i_I_* at editing rate *r*. Dividing cells pass their tapes with accumulated edits to their descendants. Editing continues until the experiment ends at time *T*. Then, cells are sampled and their barcodes are read out.

We simulated CRISPR lineage recordings along time-scaled cell phylogenies. Thus, for all tips in each tree, we obtained one vector of length *m* for non-sequential recordings and *k* vectors of length *l* for sequential recordings.

For our simulations, we fixed the experiment duration to *T* = 40. Empirical observations indicate that a few editing outcomes are much more common than others [11, 14], hence, we sampled their frequencies from an exponential distribution and scaled them to obtain scarring and insert probabilities (Table S1). We varied the remaining experimental parameters as summarized below (Table 3). We have used *Snakemake* [55] to automate the simulations.

**Table 3:** Parameters for simulating CRISPR lineage recordings.

| non-sequential |  |  | sequential |  |  |
| --- | --- | --- | --- | --- | --- |
| Parameter | Baseline | Variations | Parameter | Baseline | Variations |
| editing rate $r$ | 0.05 | 0.01, 0.15 | editing rate $r$ | 0.05 | 0.01, 0.15, 0.45 |
| number of targets $m$ | 20 | 5, 40 | number of tapes $k$ | 20 | 1, 5, 40 |
| editing height $t_e$ | 40 | 20, 30 | tape length $l$ | 5 | 2, 10, 20 |
| editing duration $\Delta t_e$ | 40 | 20 | | | |

For recordings with noise, we used versions of the CRISPR editing simulators in *TiDeTree* and *SciPhy* with two additional parameters: the silencing rate *r_s_* = 0.01, specifying the rate at which targets or tapes get lost during the editing window, and the dropout probability *p_m_* = 0.2, specifying the probability of targets or tapes being missing at sequencing at the end of the experiment. We run the simulations on larger trees (cf. Section S5.1 in Supplementary Material) and then filtered the recordings to only include cells and targets/ tapes with non-missing sites.

### 4.3 Bayesian inference

In general, Bayesian phylogenetic and phylodynamic inference [56] starts with a collection of *n* sequences, denoted alignment *A*. It assumes that the sequences evolved from a common ancestor according to a substitution model with rate matrix Q along the branches of a tree T. The tree itself resulted from a population dynamic process specified by a set of parameters *η*. The goal is to infer the unknown aspects of the past process, that is, the tree and parameters of the substitution and phylodynamic models from the sequences. Prior information on the parameters of the assumed model *M* have to be specified. Applying the Bayes rule and assumptions of independence between the components results in the expression

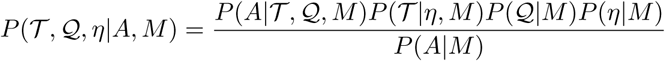

for the posterior distribution of the variables of interest. The term *P* (*A*|T, Q*, M*) is called the phylogenetic likelihood and *P* (T |*η, M*) the phylodynamic likelihood.

Due to the large dimensionality of the state space and the impractical evaluation of the marginal likelihood *P* (*A*|*M*), Monte Carlo algorithms, such as Markov chain Monte Carlo (MCMC), are commonly used to estimate the posterior distribution. The MCMC algorithm explores the state space by collecting samples from the posterior distribution. Each sample consists of the numerical parameters of the substitution and phylodynamic models, as well as the tree topology and the associated branch lengths. We employed the Bayesian MCMC framework in BEAST 2 to jointly infer time- scaled cell phylogenies and parameters of the cell population processes as well as editing process from CRISPR lineage tracing data. Our simulated barcodes constituted the input alignments.

We fit the editing (substitution) models from *TiDeTree* and *SciPhy* to the non- sequential and sequential barcodes, respectively. We inferred the editing rate and fixed the scarring and insert probabilities to true values, because we reasoned that the occurrence and relative frequency of each editing outcome can be quantified in real experiments using sequencing data.

To infer population dynamics, we used the birth-death skyline contemporary model implemented in *BDSKY* [57] for homogeneous cell populations, and the multi-type birth-death model implemented in *BDMM-Prime* [35–37] for heterogeneous cell populations. Due to identifiability reasons [33], we fixed the sampling proportion to the truth and inferred the birth (cell division), death and non-zero transition rates. Fixing the sampling proportion can also be done for empirical data as the sampling proportion at the end of the experiment can be determined.

We fixed the origin of the phylogenetic trees at 40, reflecting the experiment duration, and specified the editing window. We inferred the tree topology and branch lengths, from which we derived the tree length (the sum of all branch lengths) and tree height. For heterogeneous populations, we first reconstructed tip-typed trees [37] (meaning types ancestral to the tips were not inferred) and, subsequently, stochastically mapped ancestral type changes on the trees.

We used weakly informative distributions as priors in our analysis (Table 4). We let the analysis run for 10^8^ steps or until the effective sample size (ESS) was above 200 and discarded 10% of the analysis to account for burn-in.

**Table 4:** Prior distributions for inference.

| Parameter | Prior | Percentiles [5<sup>th</sup>, 95<sup>th</sup>] |
| --- | --- | --- |
| editing rate $r$ | LogNormal(-3, 1)<sup>1</sup> | [0.01, 0.26] |
| birth rate $\beta$ | LogNormal(-1.5, 1) | [0.04, 1.16] |
| death rate $\delta$ | Exponential(25) | [0.002, 0.12] |
| transition rate $\gamma$ | LogNormal(-3.25, 1) | [0.007, 0.20] |

In the multi-type model, we used the same prior distribution for each non-zero element of the vectors $\beta$ , $\delta$ and the matrices $\gamma$ .
<sup>1</sup>For $r = 0.45$ , we used LogNormal(-1, 1) with percentiles [0.07, 1.91].

### 4.4 Evaluation

We characterized different proxies for information content in our simulated alignments by calculating the number of different editing outcomes, the number of edits per barcode, and the pairwise Hamming distance between barcodes. We defined alignment diversity as the number of unique barcodes relative to their total number, corresponding to the proportion of sampled cells which have a unique pattern of edits accumulated across all target sites.

For each simulation and the respective inference run, and for each inferred parameter, we computed the median and the 95% highest posterior density (HPD) interval of the posterior probability distribution. To evaluate the inference performance for parameters of particular interest (editing, cell division, death, growth, and transition rates, tree height and tree length) across simulations, we calculated the following metrics:

- Coverage: fraction of times the true parameter was contained in HPD interval.
- Relative bias: difference between the posterior median and the true parameter, divided by the truth.
- Relative absolute error: absolute difference between the posterior median and the true parameter, divided by the truth.
- Relative HPD width: difference between the upper and lower bound of the posterior 95% HPD interval divided by the true value of the parameter.
- HPD proportion (only for parameters with an explicitly specified prior distribution; but not for tree height and tree length): difference between the upper and lower bound of the posterior 95% HPD interval divided by the difference between the upper and lower bound of the 95% credible interval of the prior distribution.

For synchronous trees with birth only, the true death rate was 0, so we normalized the metrics by the mean of the median estimates instead of the truth. To be able to compare the inference of the death rate across population models, we also calculated not normalized metrics for this parameter.

To evaluate the reconstruction of cell phylogenies, we derived for each inference the 95% credible set of tree topologies, that is the smallest set of all tree topologies that accounts for 95% of the posterior probability. We measured its relative size by dividing the number of trees in the credible set by the number of sampled trees. Then, we calculated tree coverage as the fraction of times the true tree was in the credible set across simulations. Furthermore, we summarized the posterior trees to the maximum clade credibility (MCC) tree with the node heights rescaled to the posterior median node heights for the clades contained in the tree. We quantified the (dis)similarity between each MCC (‘inferred’) tree and the corresponding true phylogenetic tree in three ways. We computed the weighted Robinson-Foulds (RF) distance [38, 39] which considers branch lengths alongside tree topology. Additionally, we computed two generalized topological RF distance metrics, Nye similarity [58] and Shared Phylogenetic Information [59], which account for differences between similar but not identical pairs of tree splits. We normalized the metrics by their maximum values so that they fall between 0 and 1. These calculations were done using the R packages *TreeDist* [60] and *phangorn* [61]. Additionally, we computed the Wasserstein and Kolmorogov–Smirnov distance between the branch length distributions of the inferred and true trees.

For multi-type phylogenies, we evaluated the inference of ancestral cell types by identifying clades that a given MCC tree and the corresponding true tree have in common, and calculating the proportion of internal nodes in these clades with correctly assigned types.

Additionally, we used the R packages *treeio* [62, 63], *tracerer* [64] and *tidyverse* [65] for evaluation.

### 4.5 Approximation

As described in 4.3, we fitted the birth-death model, parametrized by a birth (cell division) and death rate, to all homogeneous cell phylogenies. However, the population model underlying synchronous trees is parametrized differently, by the number of time points at which all living cells simultaneously divide, and optionally, the probability of cell death. To be able to evaluate the phylodynamic inference for phylogenetic trees with synchronous cell divisions, we approximated their ‘generative’ birth and death rates in two ways. We compared these approximate ‘generative’ birth and death rates to the estimated birth and death rates.

#### Approximation per lineage

First, we consider the time intervals between subsequent cell divisions and deaths. In the birth-death model, both events follow a Poisson process, so the expected waiting time for the next birth event is exponentially distributed with parameter *β*, and the expected waiting time for the next death event is exponentially distributed with parameter *δ*. The expected waiting time for the next event of any kind is exponential with parameter *β* + *δ* and mean 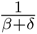. Then, a cell divides with probability 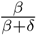 or dies with probability 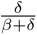.

In our synchronous tree simulations, cell divisions and deaths occur after fixed time intervals. We now approximate this process with ‘generative’ birth and death rates. If *n* denotes the number of cell division time points within [0*, T*], the time interval between subsequent divisions is 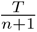. Taking the inverse results in a per-lineage event rate of 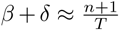. Given a death probability *p_d_* and noting that *δ* = *p_d_* · (*β* + *δ*), the birth and death rates are approximated by the ‘generative’ rates

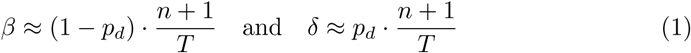

In birth-only synchronous trees, *p_d_* = 0, and thus the rates are simply 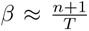 and *δ* = 0.

#### Approximation on population level

In our second approximation, we derive the birth and death rates from the expected number of cells at the end of the tree generating process, at time *T*. Under the birth-death model, the expected number of living individuals (cells) after arbitrary time *t* is

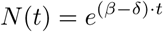

assuming that the population consists initially of one cell [32]. Hence, the number of cells grows exponentially through time as long as *β > δ*. Given a sampling probability *ρ*, on average *ρ* · *e*^(*β−δ*)*·T*^ cells are sampled at time *T*.

Under synchronous cell divisions and sampling fraction *ρ*, the expected number of cells after *n* division time points is *ρ* · 2*^n^*. Hence, for birth-only trees, in which *δ* = 0, we can approximate

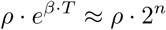

meaning the ‘generative’ birth rate is 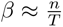 ln 2.

In synchronous trees with cell death, instead of considering the expected number of surviving cell lineages, we used the event rate *β* +*δ* to compare the expected number of cells *ρ* · 2*^n^*to the birth-death model, treating deaths as birth events in the calculation.

We arrive at the equation

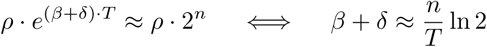

For a given death probability *p_d_*, approximated birth and death rates on the population level are thus

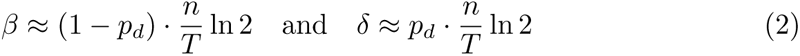

Again, for synchronous trees with birth only, 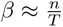 ln 2 and *δ* = 0. We use these approximations as the ‘generative’ birth and death rates.

Altogether, for *n* → ∞, the population level birth and death rates differ by a factor of ln 2 from the per lineage rates. We suggest that the difference stems from the assumed base in the population growth functions. The per lineage rates would match with the population level rates, if the population growth over time was exponential with base *e*, as in the birth-death model. However, when cell divisions are synchronous, the population growth over time is slower - exponential with base 2. Multiplying the per lineage rates with factor ln 2 changes the base in the growth function from *e* to 2. Applying the inference framework to our simulated data, we have observed that the posterior medians consistently approached the birth and death rates approximated on the population level. Therefore, we treated those as the ‘true’ rates in the evaluation. Our results further imply that birth rates inferred under the birth-death model can be divided by ln 2 to estimate the doubling time of cell populations growing by synchronous and regular cell divisions in empirical analyses.

## Data and Code Availability

Code to rerun the simulations and analysis and generate the figures is available at https://github.com/pilarskj/celldev.

## Author contributions

**Julia Pilarski**: Conceptualization, Data curation, Formal analysis, Investigation, Methodology, Project Administration, Software, Validation, Visualization, Writing – original draft, Writing – review & editing. **Tanja Stadler**: Conceptualization, Formal analysis, Funding Acquisition, Investigation, Methodology, Resources, Supervision, Writing – review & editing. **Sophie Seidel**: Conceptualization, Formal analysis, Investigation, Methodology, Resources, Supervision, Writing – review & editing.

## Acknowledgments

The authors thank Antoine Zwaans, Nicola Mulberry and other members of the cEvo group at ETH Zurich for helpful discussions, and Jay Shendure and Florence Chardon for helpful comments on the manuscript.

## Supporting Information

The supplementary material contains Tables S1-S3 and Figures S1-S17 organized into the following sections:

**S1** Simulation of barcodes

**S2** The baseline

**S3** Varying experimental parameters

**S4** Evaluating sequential editing

**S5** Increasing the sample size and subsampling cells

**S5.1** Simulation of large recordings

**S5.2** Performance of tree and parameter inference

**S5.3** Runtime and convergence

**S6** Filtering out noisy data

**S7** Inferring cell differentiation dynamics

