## Supplementary material for "Assessing the inference of single-cell phylogenies and population dynamics from CRISPR lineage recordings"

#### S1 Simulation of barcodes

In Table S1, we report the probabilities of editing outcomes sampled and used for simulating CRISPR lineage recordings (cf. Section 4.2).

#### S2 The baseline

Below, we provide additional results for simulations in homogeneous cell populations at baseline experimental parameters (as introduced in Section 2.2). Fig. S1 shows the posterior distributions of parameters inferred from each simulated recording. Fig. S2 depicts example phylogenetic trees and their posteriors. Fig. S3 provides additional metrics for phylogenetic reconstruction across tree generating processes. Fig. S4 compares the inferred and true branch length distributions for the different population-dynamic models. Fig. S5 shows that improved phylogenetic reconstruction is associated with reduced uncertainty in the branching times, which propagates to reduced uncertainty in the phylodynamic estimates.

**Table S1**: Edit probabilities for simulating CRISPR lineage recordings

| Index $k$ | Scarring probability $s_k$ | Insert probability $i_k$ |
|---|---|---|
| 1 | 0.0021 | 0.0109 |
| 2 | 0.0078 | 0.0266 |
| 3 | 0.0812 | 0.0690 |
| 4 | 0.0105 | 0.0018 |
| 5 | 0.0039 | 0.0707 |
| 6 | 0.0309 | 0.0803 |
| 7 | 0.0025 | 0.2657 |
| 8 | 0.0513 | 0.0038 |
| 9 | 0.0021 | 0.0445 |
| 10 | 0.0731 | 0.0106 |
| 11 | 0.0441 | 0.1265 |
| 12 | 0.0038 | 0.1935 |
| 13 | 0.0195 | 0.0961 |
| 14 | 0.0455 | |
| 15 | 0.0072 | |
| 16 | 0.0210 | |
| 17 | 0.0136 | |
| 18 | 0.0116 | |
| 19 | 0.0065 | |
| 20 | 0.0014 | |
| 21 | 0.0225 | |
| 22 | 0.0056 | |
| 23 | 0.0067 | |
| 24 | 0.0370 | |
| 25 | 0.0189 | |
| 26 | 0.0281 | |
| 27 | 0.0070 | |
| 28 | 0.0132 | |
| 29 | 0.0002 | |
| 30 | 0.0099 | |
| 31 | 0.0397 | |
| 32 | 0.0190 | |
| 33 | 0.0099 | |
| 34 | 0.0017 | |
| 35 | 0.0160 | |
| 36 | 0.0359 | |
| 37 | 0.0001 | |
| 38 | 0.0063 | |
| 39 | 0.0006 | |
| 40 | 0.0494 | |
| 41 | 0.0396 | |
| 42 | 0.0027 | |
| 43 | 0.0148 | |
| 44 | 0.0391 | |
| 45 | 0.0421 | |
| 46 | 0.0360 | |
| 47 | 0.0032 | |
| 48 | 0.0242 | |
| 49 | 0.0004 | |
| 50 | 0.0306 | |

A: non-sequential recordings

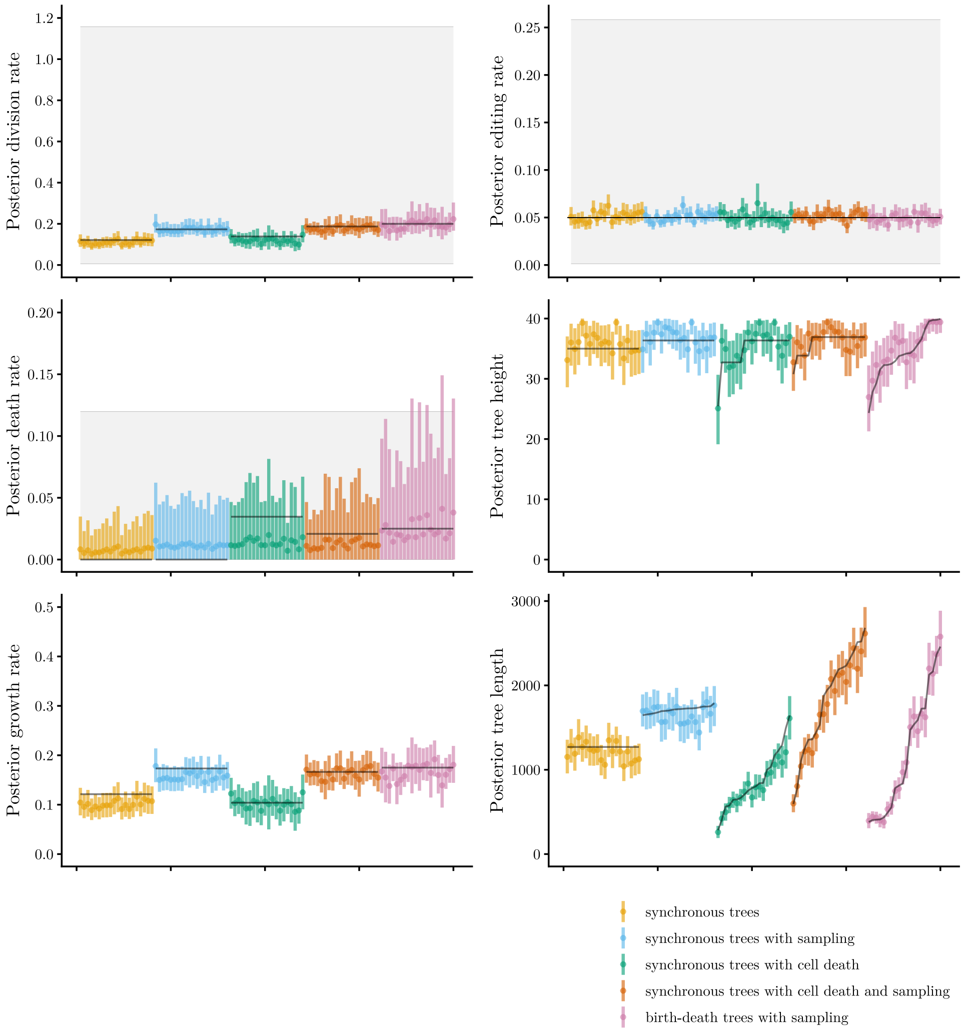


**Fig. S1**: Parameters inferred from (A) non-sequential and (B) sequential CRISPR lineage recordings at baseline. In each graph, points indicate the medians and bars show the 95% HPD intervals of the parameter estimates per simulation (x-axis). Colors represent the different tree generating processes. Black lines display the true parameters. Grey shaded areas represent the 95% HPD intervals of prior distributions.

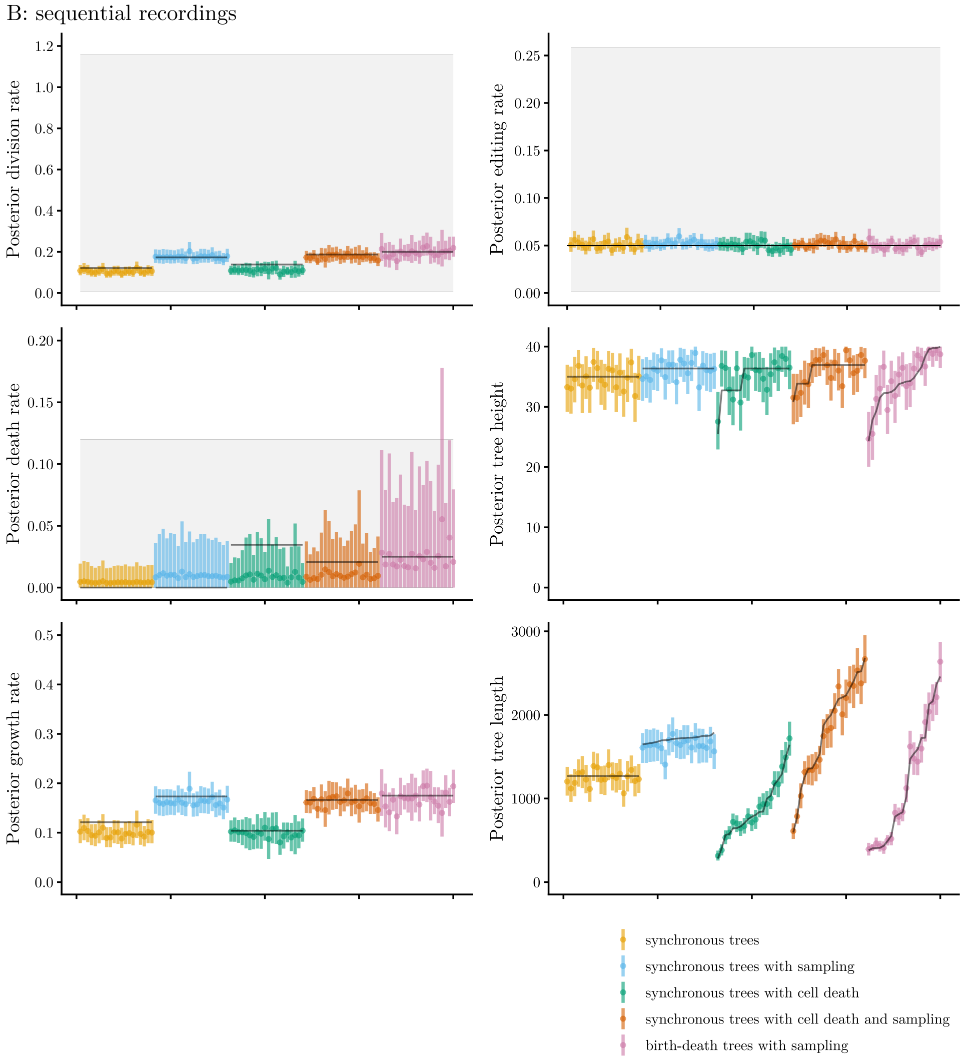


**Fig. S1**: Parameters inferred from (A) non-sequential and (B) sequential CRISPR lineage recordings at baseline. (cont.)

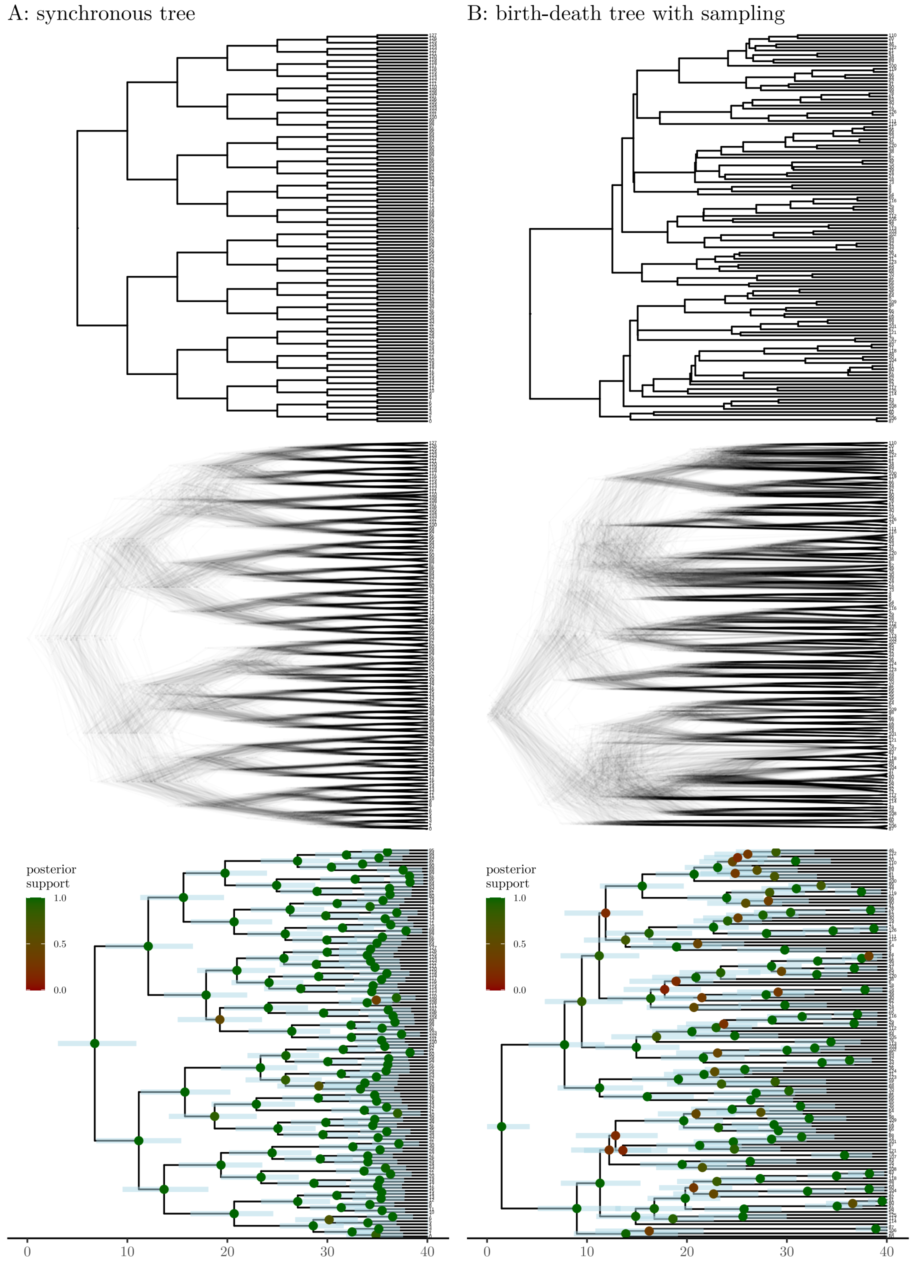


**Fig. S2**: Example (A) synchronous tree and (B) birth-death tree with sampling. The panels show from top to bottom: the true tree, a subset of the posterior tree set, the maximum clade credibility (MCC) tree inferred from the corresponding simulated sequential recording (at baseline). In the MCC tree, light blue bars indicate the 95% HPD intervals around internal node heights. Internal nodes are colored by the posterior support of the tree split. The trees are time-scaled.

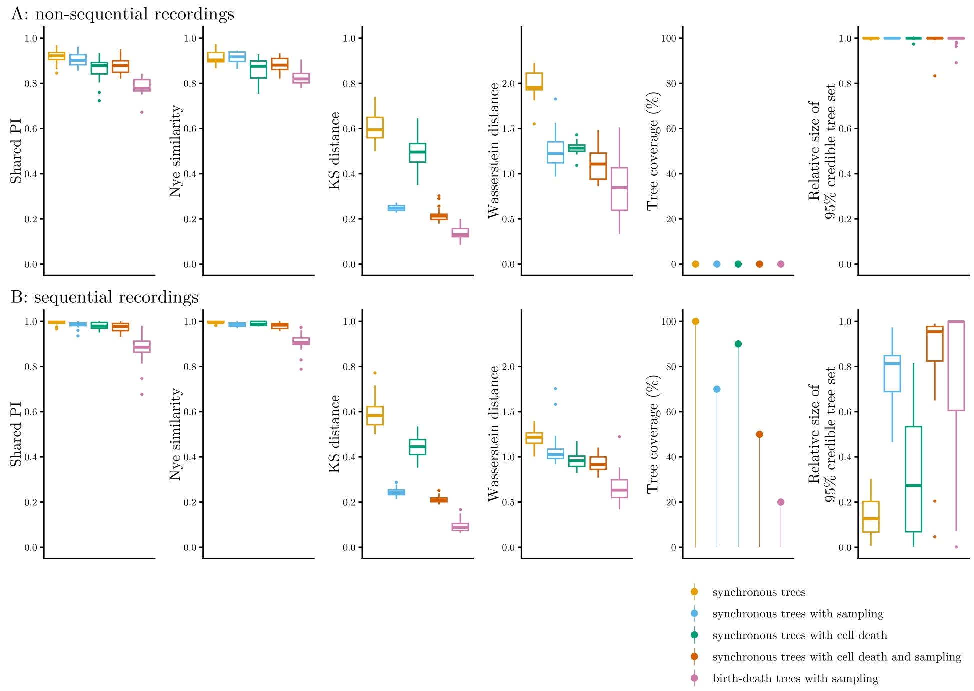


**Fig. S3**: Additional metrics for assessing phylogeny reconstruction from baseline (A) non-sequential and (B) sequential CRISPR lineage recordings. The graphs show from left to right: Shared Phylogenetic Information (PI) and Nye similarity between the inferred and true trees, the Kolmogorov-Smirnov (KS) and Wasserstein distance between branch length distributions, the tree coverage, and the relative size of the 95% credible tree set. The metrics are summarized across 20 simulations per tree generating process (indicated by colors). Dashed lines display thresholds to facilitate visual comparability.

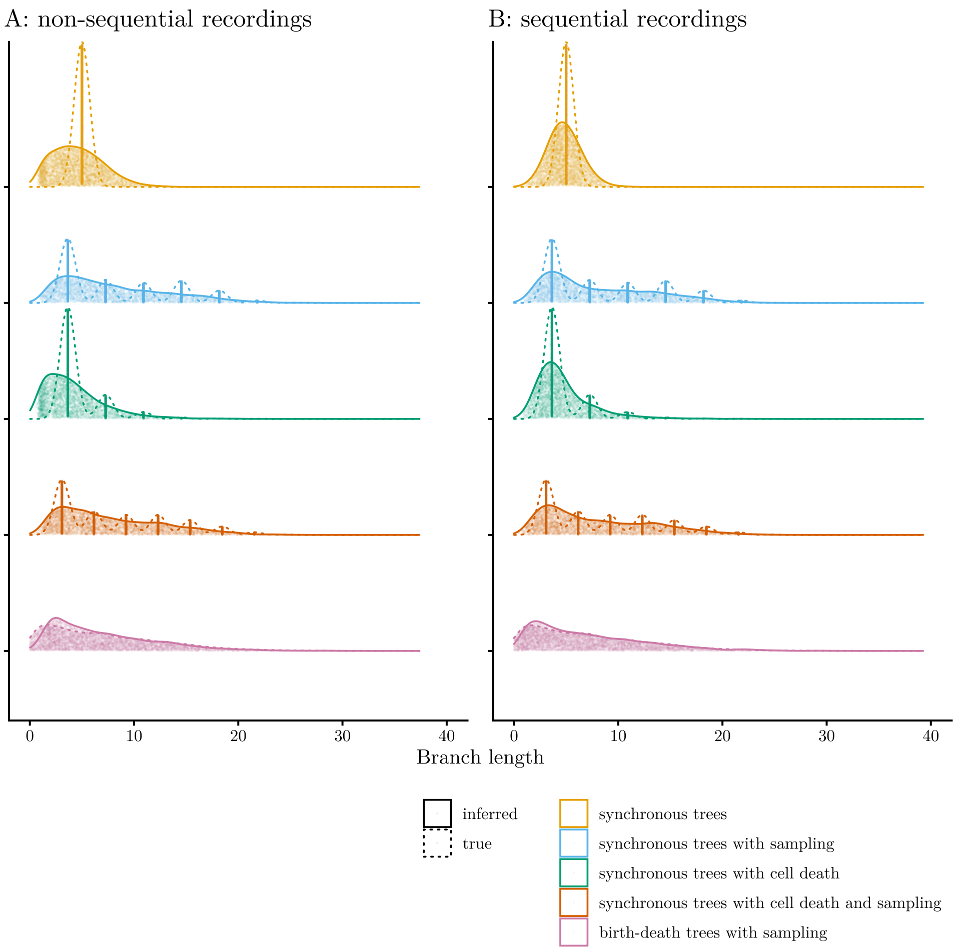


**Fig. S4**: Inferred versus true branch length distributions from baseline (A) non-sequential and (B) sequential CRISPR lineage recordings. Lines indicate smoothed density estimates for the true (solid) and inferred (dashed) branch length distributions. Light points show the individual branch lengths, aggregated across 20 trees per tree generating process (indicated by colors).

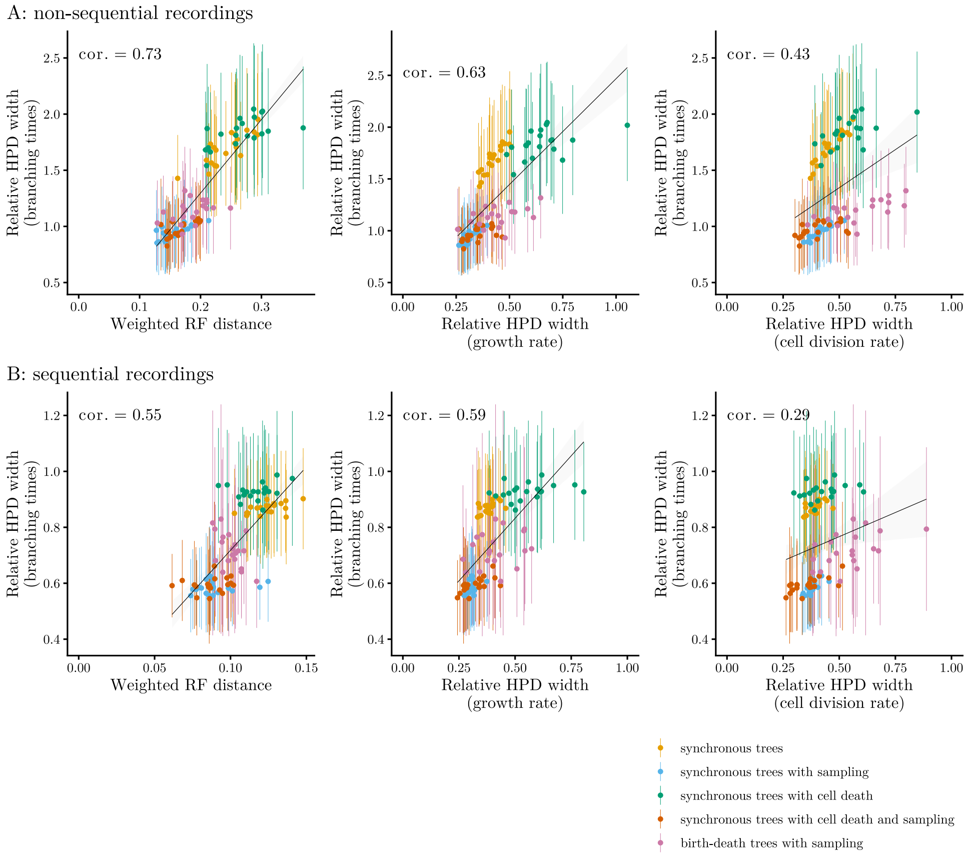


**Fig. S5**: Associations for (A) non-sequential and (B) sequential CRISPR lineage recordings at baseline. Graphs show correlations of the weighted RF distance between the inferred and true trees (each point corresponds to one simulated tree), uncertainty in the branching times (as measured by the relative width of 95% HPD posterior intervals around the internal node heights – pointranges indicate the mean ± standard deviation/2 across internal nodes in the tree), and uncertainty in the phylodynamic estimates (as measured by the relative HPD widths of posterior growth and cell divisions rates per simulation). The different tree generating processes are indicated by colors. Black lines indicate the linear trends (least squares fit). cor.: Kendall's $\tau$ correlation coefficient.

### S3 Varying experimental parameters

Below, we show detailed results for simulations in homogeneous cell populations at varying experimental parameters (as described in Section 2.3). In Table S2, we report the relative number of edited sites in simulated CRISPR lineage recordings under different settings. In Figs. S6-S7, we compare parameter and tree inference for all scenarios across tree generating processes. Additionally, in Fig. S8, we visualize the improvements in phylogenetic and phylodynamic inference as a function of barcode diversity.

**Table S2**: Recorder saturation at varying experimental parameters

| Experimental design | | Recorder | |
|---|---|---|---|
| | | non-sequential | sequential |
| baseline | | 17/20 (85%) | 39.5/100 (39.5%) |
| editing rate | 0.01 | 6/20 (30%) | 8/100 (8%) |
| | 0.15 | 20/20 (100%) | 90/100 (90%) |
| number of targets/ tapes | 5 | 4/5 (80%) | 10/25 (40%) |
| | 40 | 34/40 (85%) | 79/200 (39.5%) |
| tape length | 2 | | 29/40 (72.5%) |
| | 10 | | 39/200 (19.5%) |
| editing window | first half | 12/20 (60%) | |
| | mid | 13/20 (65%) | |
| | second half | 13/20 (65%) | |

The entries indicate the median number of edited sites relative to the number of available sites across all cells per recording, summarized across 100 simulations per setting.

A: non-sequential recordings

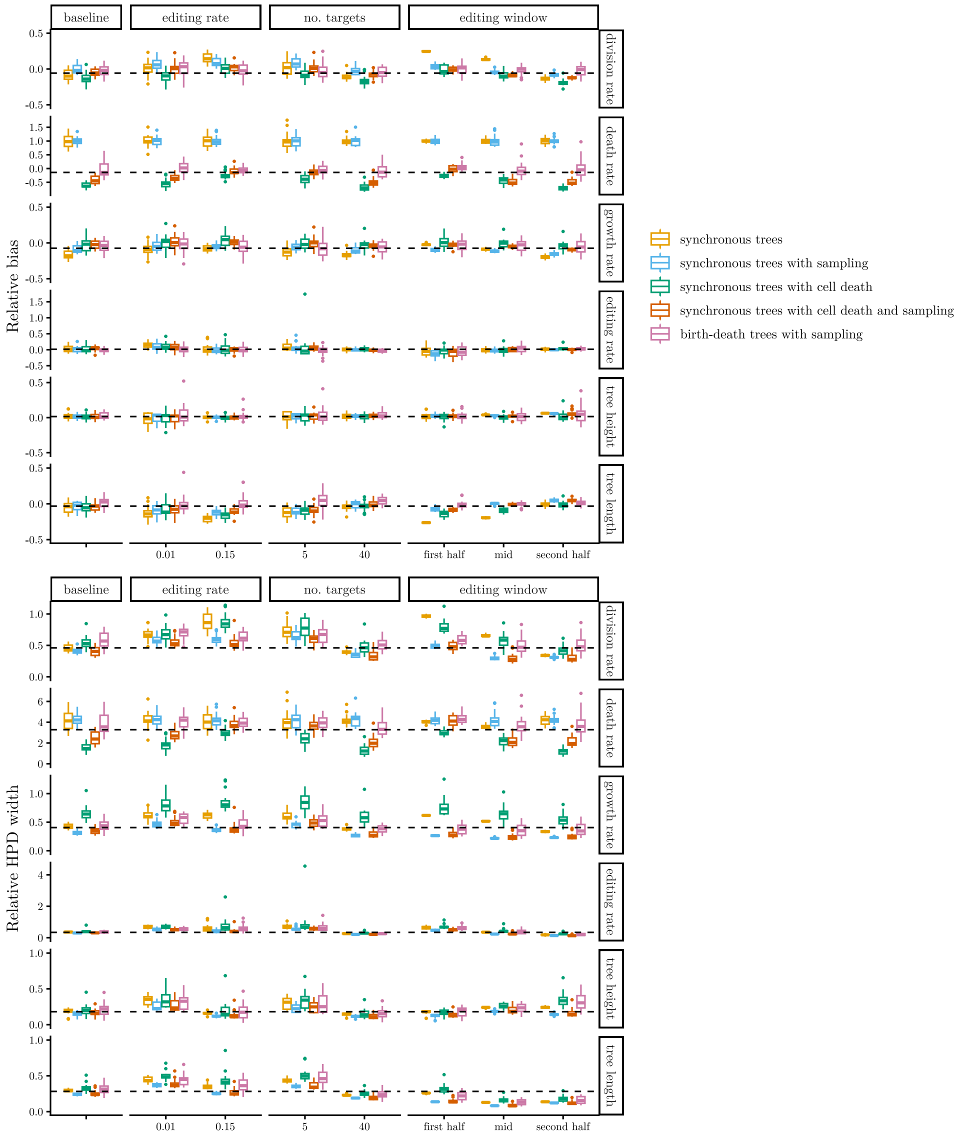


**Fig. S6**: Parameter inference performance from (A) non-sequential and (B) sequential CRISPR lineage recordings with varying experimental parameters. Vertical facets indicate the varied experimental parameters, horizontal facets indicate the inferred parameters. At baseline, the editing rate was 0.05, the recorders carried 20 targets or 20 tapes of length 5, respectively, and editing occurred throughout the entire experiment. Dashed lines display the baseline medians (across trees). The graphs show the relative bias (top) and relative HPD width (bottom) of the inferred parameters. The metrics are summarized across 20 simulations per tree generating process (indicated by colors) per setting.

B: sequential recordings

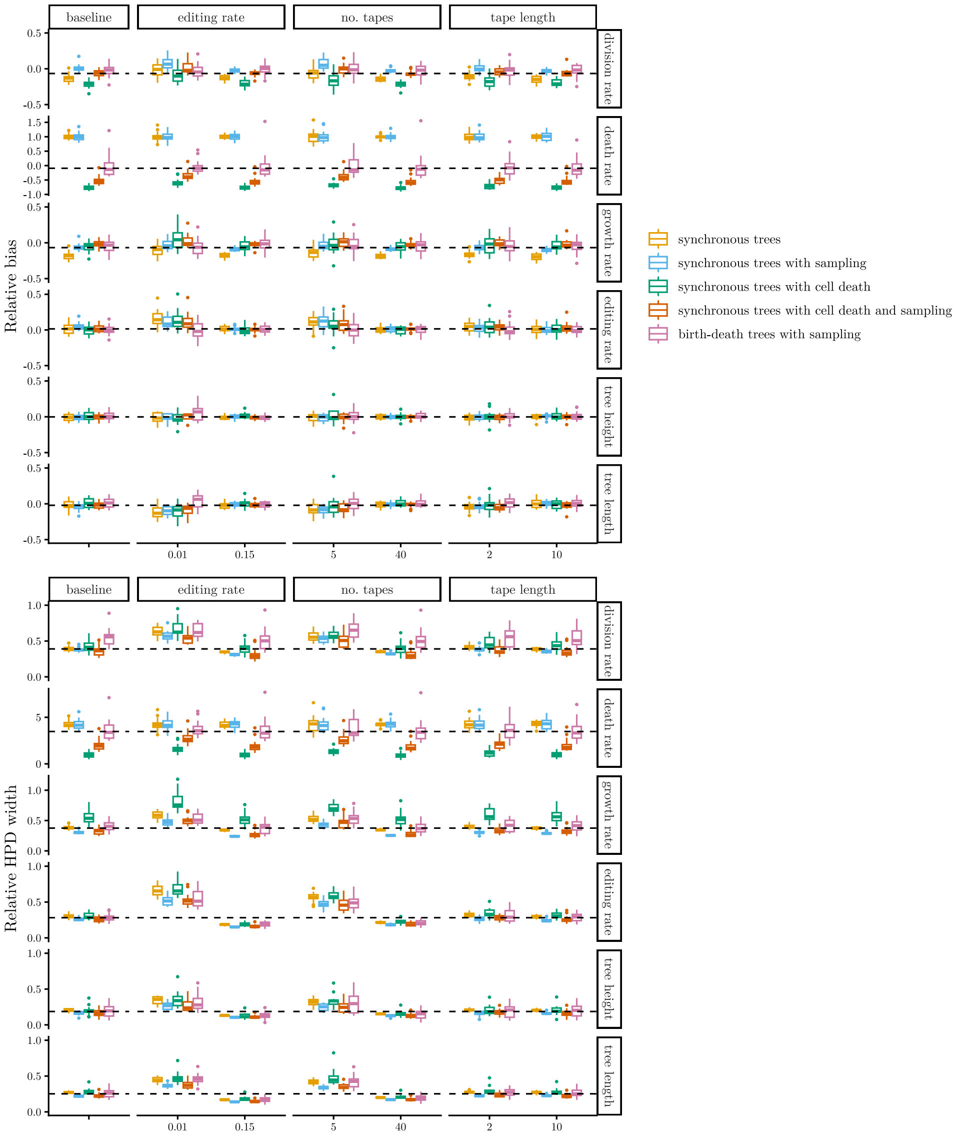


**Fig. S6**: Parameter inference performance from (A) non-sequential and (B) sequential CRISPR lineage recordings with varying experimental parameters. (cont.)

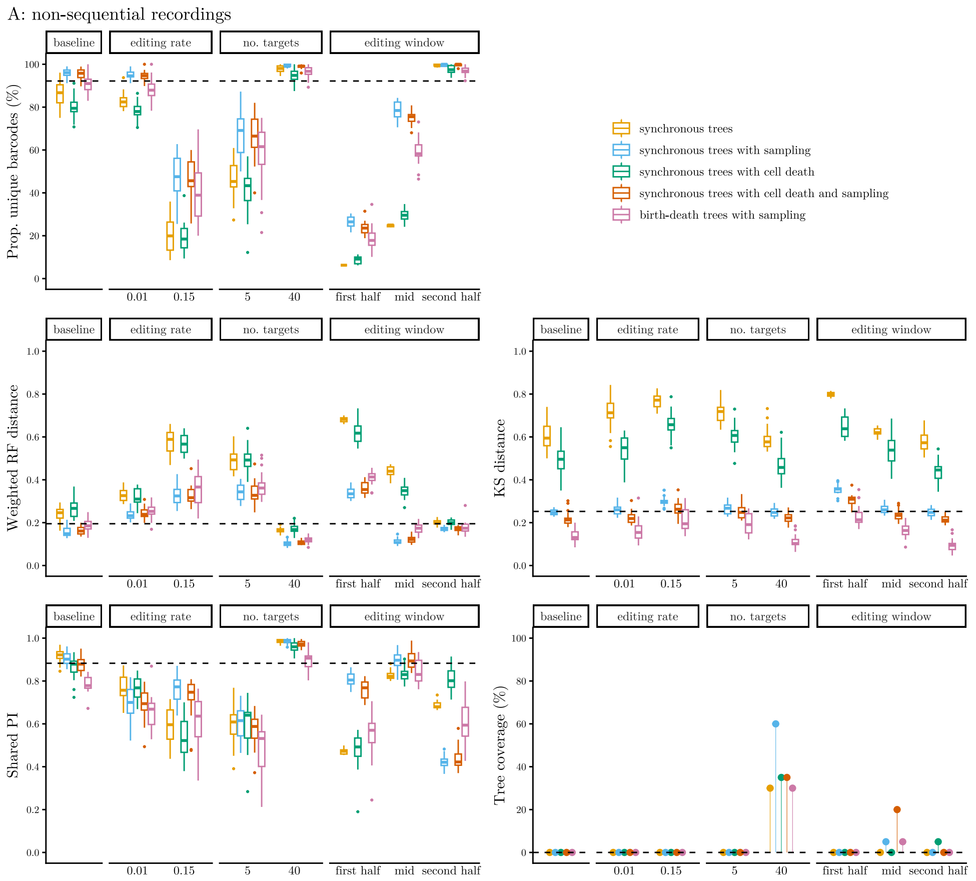


**Fig. S7**: Additional metrics for assessing phylogeny reconstruction from (A) non-sequential and (B) sequential CRISPR lineage recordings with varying experimental parameters. Facets indicate the varied experimental parameters. At baseline, the editing rate was 0.05, the recorders carried 20 targets or 20 tapes of length 5, respectively, and editing occurred throughout the entire experiment. Dashed lines display the baseline medians (across trees). The top panels summarize barcode diversity per recording, defined as the proportion of unique barcodes among the sampled cells. The graphs below show the different tree metrics. The results are summarized across 20 simulations per tree generating process (indicated by colors) per setting.

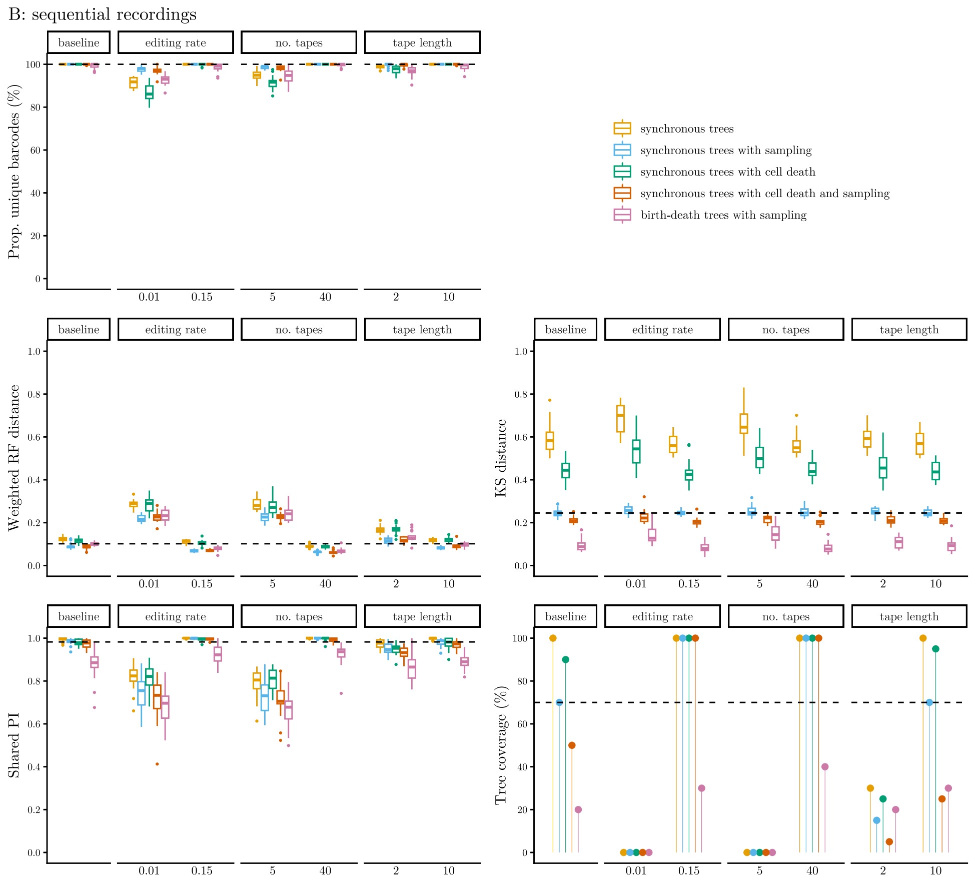


**Fig. S7**: Additional metrics for assessing phylogeny reconstruction from (A) non-sequential and (B) sequential CRISPR lineage recordings with varying experimental parameters. (cont.)

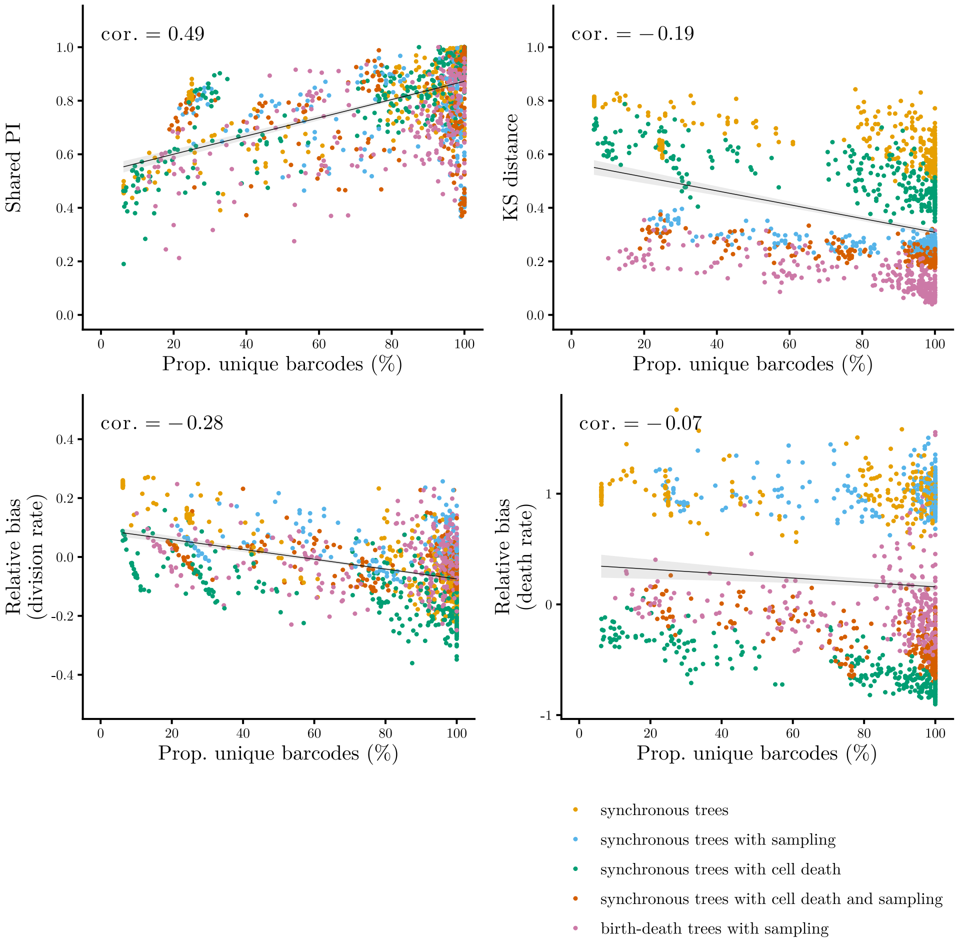


**Fig. S8**: Associations between barcode diversity (measured by the proportion of unique barcodes per recording), accuracy of phylogenetic reconstruction in terms of tree topology (measured by the shared PI between the inferred and true trees) and in terms of branch lengths (measured by the KS distance between branch length distributions), and accuracy of phylodynamic inference (evaluated with the relative bias of the cell division and death rate estimates) across experimental scenarios and tree generating processes (indicated by colors). Black lines indicate the linear trends (least squares fit). cor.: Kendall's $\tau$ correlation coefficient.

#### S4 Evaluating sequential editing

In Fig. S9, we compare the sequential to non-sequential accumulation of edits in CRISPR lineage recorders across population-dynamic models (as summarized in Section 2.4).

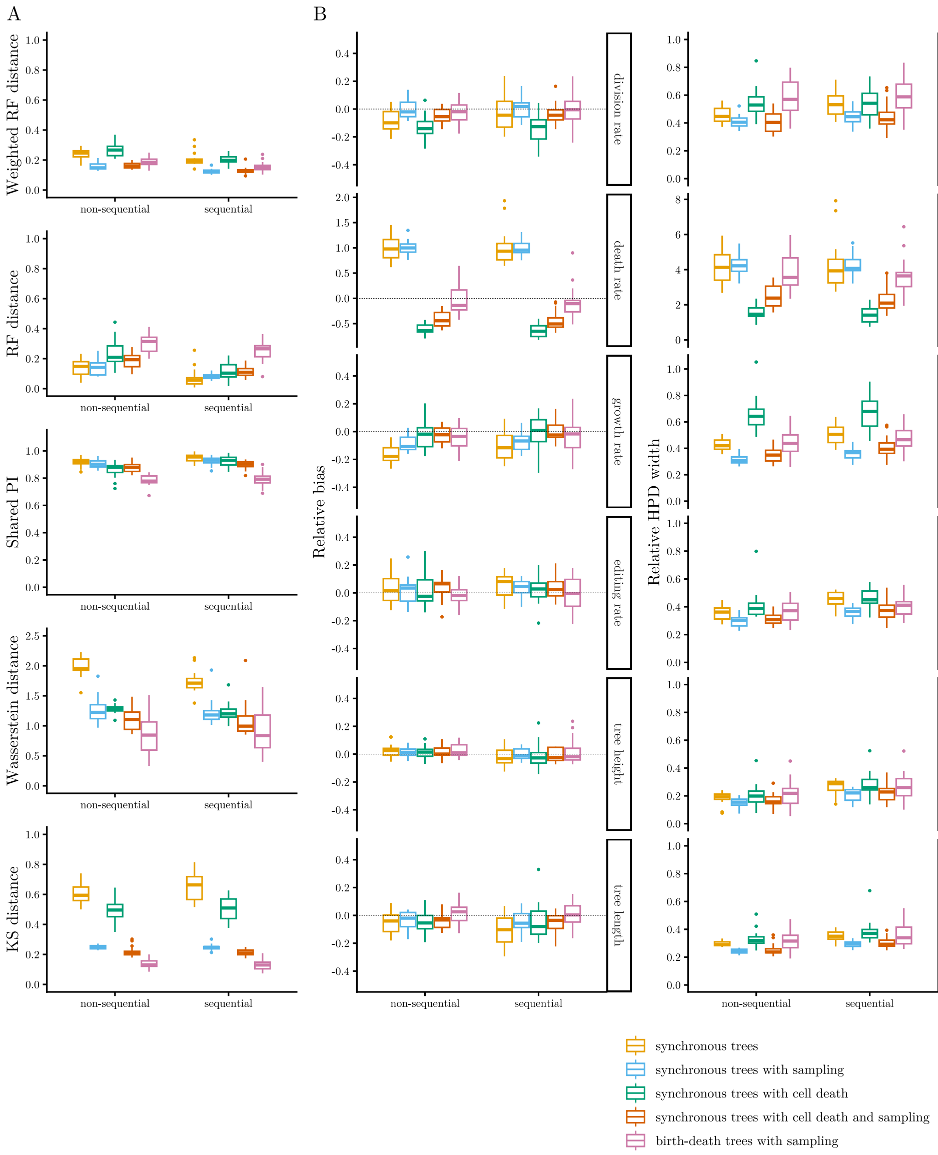


**Fig. S9**: Comparison of the sequential and non-sequential accumulation of edits in CRISPR recorders. Panel (A) shows from top to bottom: the weighted RF distance, topological RF distance, and shared PI between the inferred and true trees, and the Wasserstein and KS distance between branch length distributions. Panel (B) shows the relative bias (left) and relative HPD width (right) of the inferred parameters. The results are summarized across 20 simulations per tree generating process (indicated by colors) per setting.

### S5 Increasing the sample size and subsampling cells

#### S5.1 Simulation of large recordings

We performed additional simulations of CRISPR lineage recordings on tenfold larger trees with roughly 1000 tips to assess how the sample size affects the inference, and to test whether our results are generalizable to larger samples. First, we generated 20 trees for the five population dynamics, mimicking the development of homogeneous cell populations. As before, we started each tree generating process with a single cell at time $t = 0$ and let the cell populations evolve for 40 time units. To reach a tree size of 200 to 2000 tips, we used parameters as summarized in Table S3. For simulating birth-death trees with sampling efficiently, we used the BEAST 2 [1] package ReMASTER [2].

**Table S3**: Parameters for simulating large homogeneous phylogenetic trees

| Trees | $n$ | $p_d$ | $\beta$ | $\delta$ | $\beta - \delta$ | $\rho$ |
|---|---|---|---|---|---|---|
| synchronous | **10** | **0** | 0.17 | 0 | 0.17 | **1** |
| synchronous with sampling | **13** | **0** | 0.225 | 0 | 0.225 | **0.1** |
| synchronous with cell death | **13** | **0.15** | 0.19 | 0.03 | 0.16 | **1** |
| synchronous with cell death and sampling | **16** | **0.12** | 0.24 | 0.03 | 0.17 | **0.1** |
| birth-death with sampling | - | 0.115 | **0.27** | **0.035** | 0.235 | **0.1** |

We specified the values in bold, and approximated the remaining parameters (on population level).

Second, we simulated non-sequential and sequential CRISPR lineage recordings along the trees using *TiDeTree* and *SciPhy*, respectively, with baseline experimental parameters (cf. Table 3). Third, we run Bayesian phylogenetic and phylodynamic inference to reconstruct the time-scaled lineage trees and estimate the editing, cell division, and death rates from simulated barcodes. We followed the procedure described in Sections 4.3 and 4.4.

Next – for the purpose of a fair comparison of the population-dynamic parameters, and to directly assess the effects of subsampling data for analysis – we downsampled the large trees with a sampling proportion of 10% to contain only 1% of tips. Again, we simulated recordings along them, and inferred lineage trees and population-dynamic parameters from such sparser samples. In this case, we set $\rho = 0.01$ for phylodynamic inference under the birth-death model with sampling.

#### S5.2 Performance of tree and parameter inference

Below, we compare the outputs of tree and parameter inference from large recordings at 10% sampling to smaller recordings at 1% sampling.

Consistent with Bayesian theory, posterior distributions of the inferred parameters shrunk stronger relative to prior distributions at higher sample size (Fig. S10). Consequently, biases in the cell division and death rates due to phylodynamic model misspecification became more pronounced. As the posteriors became relatively narrower, they concentrated around biased estimates and failed to capture the true values in more simulations. This is especially visible for death rates in the case of synchronous trees.

Overall, the analysis on large recordings yielded concordant trends with those reported in Section 2.2 – considering the accuracy of phylogenetic reconstruction and population-dynamic parameter inference, both across tree generating processes and the two lineage recorders (Fig. S11). As before, the weighted RF distance between the inferred and true trees, normalized by tree lengths, oscillated around 0.2 for non-sequential recordings and around 0.1 for sequential recordings. Similar accuracy was achieved at sparser sampling. These results suggest that our findings across the various experimental scenarios should hold when more cells are sequenced, and the lineage recordings are downsampled for analysis.

A: non-sequential recordings

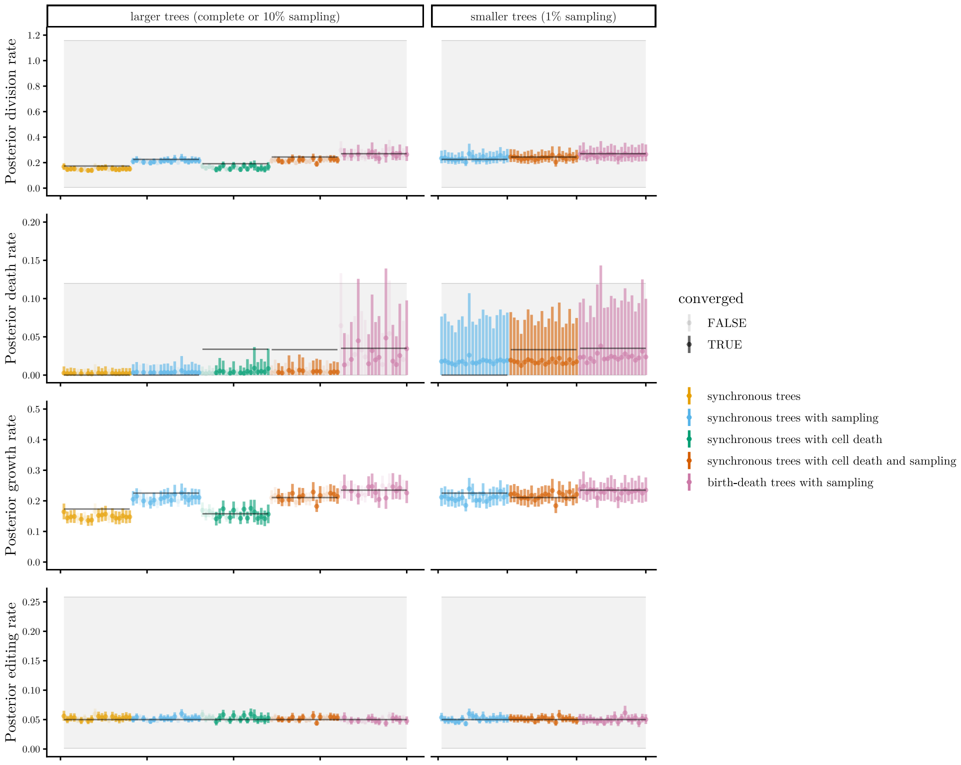


**Fig. S10**: Parameters inferred from (A) non-sequential and (B) sequential CRISPR lineage recordings simulated along large (left) and downsampled (right) trees. In each graph, points indicate the medians and bars show the 95% HPD intervals of the parameter estimates per simulation (x-axis). Colors represent the different tree generating processes. Black lines display the true parameters. Grey shaded areas represent the 95% HPD intervals of prior distributions.

B: sequential recordings

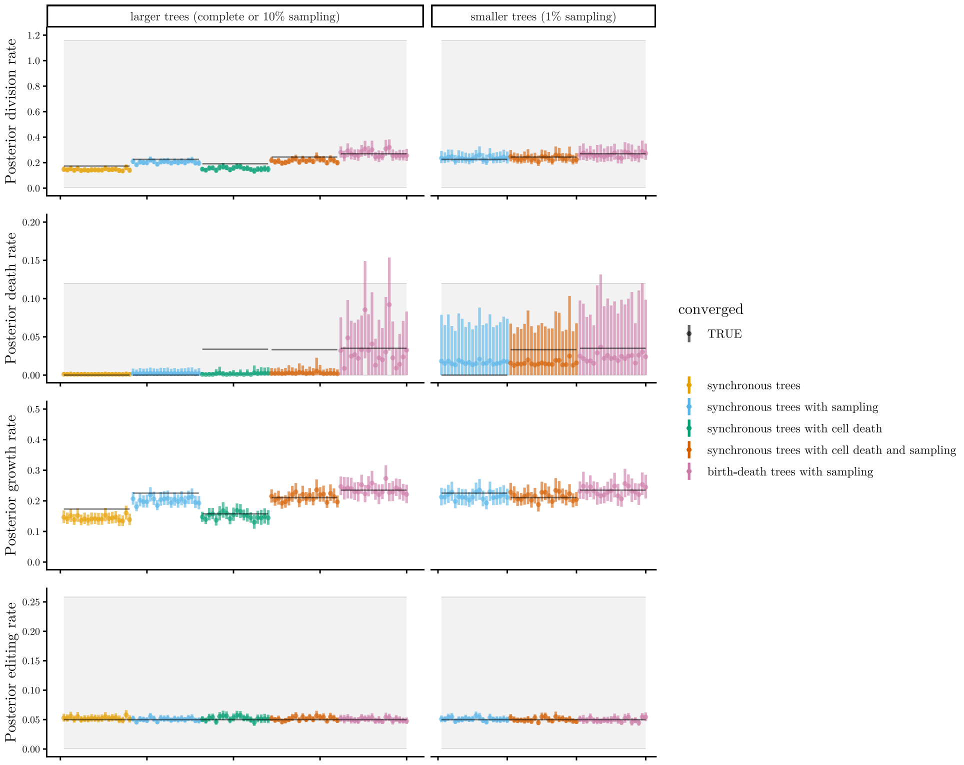


**Fig. S10**: Parameters inferred from (A) non-sequential and (B) sequential CRISPR lineage recordings simulated along large (left) and downsampled (right) trees. (cont.)

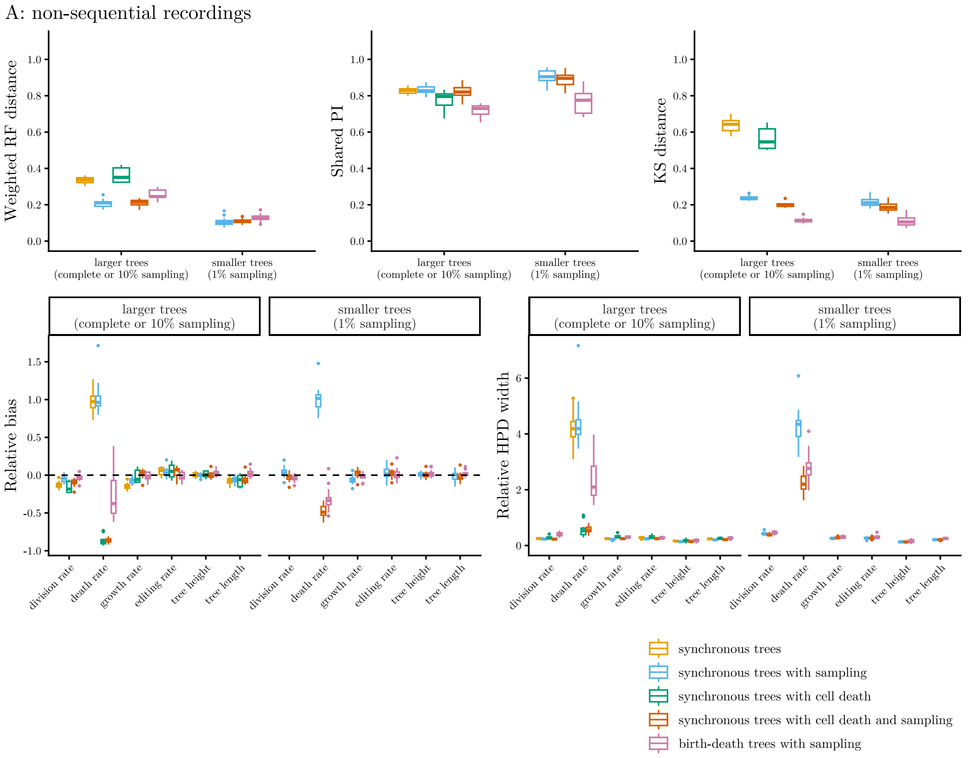


**Fig. S11**: Tree and parameter inference performance from (A) non-sequential and (B) sequential CRISPR lineage recordings simulated along large and downsampled trees. Panels (A) show from left to right: the weighted RF distance and shared PI between the inferred and true trees, and the KS distance between branch length distributions. Panels (B) show the relative bias (left) and relative HPD width (right) of the inferred parameters. The results are summarized across simulations per tree generating process (indicated by colors), for which the inference MCMC runs converged.

B: sequential recordings

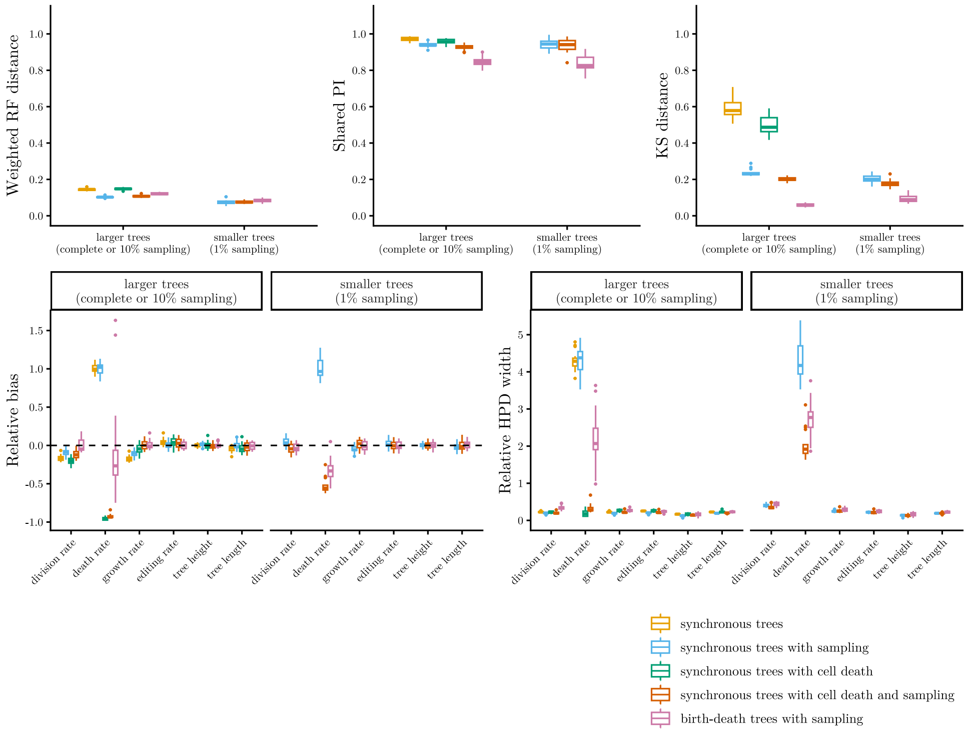


**Fig. S11**: Tree and parameter inference performance from (A) non-sequential and (B) sequential CRISPR lineage recordings simulated along large and downsampled trees. (cont.)

##### S5.3 Runtime and convergence

Currently, the Bayesian methods are limited to reconstruct and analyze trees with only few hundreds or thousands tips. There are multiple reasons for such poor scalability: First, at each iteration of an MCMC (Markov chain Monte Carlo) run, the phylogenetic likelihood is recomputed (cf. Section 4.3). In the BEAST2 packages *TiDeTree* [3] and *SciPhy* [4], this step involves Felsenstein's pruning [5] – an algorithm that recursively traverses the tree and involves dynamic programming to calculate likelihoods over subtrees. The runtime of this algorithm is linear in the number of tips. Second, the tree space grows super-exponentially with tree size, making exhaustive exploration infeasible for samples with $\gg$ 1K cells. Overall, at increasing sample size, the MCMC runs have a both a higher runtime per iteration, and require more iterations – involving tree and parameter moves – to converge.

In practice, inference on large simulated recordings required several days to approach convergence (Fig. S12A). We terminated the MCMCs after 480h, at which point only 70 out of 100 runs on non-sequential recordings had converged, reaching a posterior effective sample size (ESS) of at least 200. We estimated that some MCMC runs would require almost three times as long to converge. By contrast, all runs on sequential recordings converged within 100h.

In general, inference under *SciPhy* is multifold faster than under *TiDeTree* (Fig. S12B). The primary reason is an optimization within the implementation: in *SciPhy*, subtree likelihoods are cached and restored for any partitions unaffected by a given tree move. On the large simulated recordings, *SciPhy* was on average four times faster than *TiDeTree* in terms of runtime per million MCMC samples. Interestingly, this performance gap doubled when accounting for the total time until convergence. The additional speed-up likely stems from the increased information content in sequential recordings, representing an inherent feature of the simulated data.

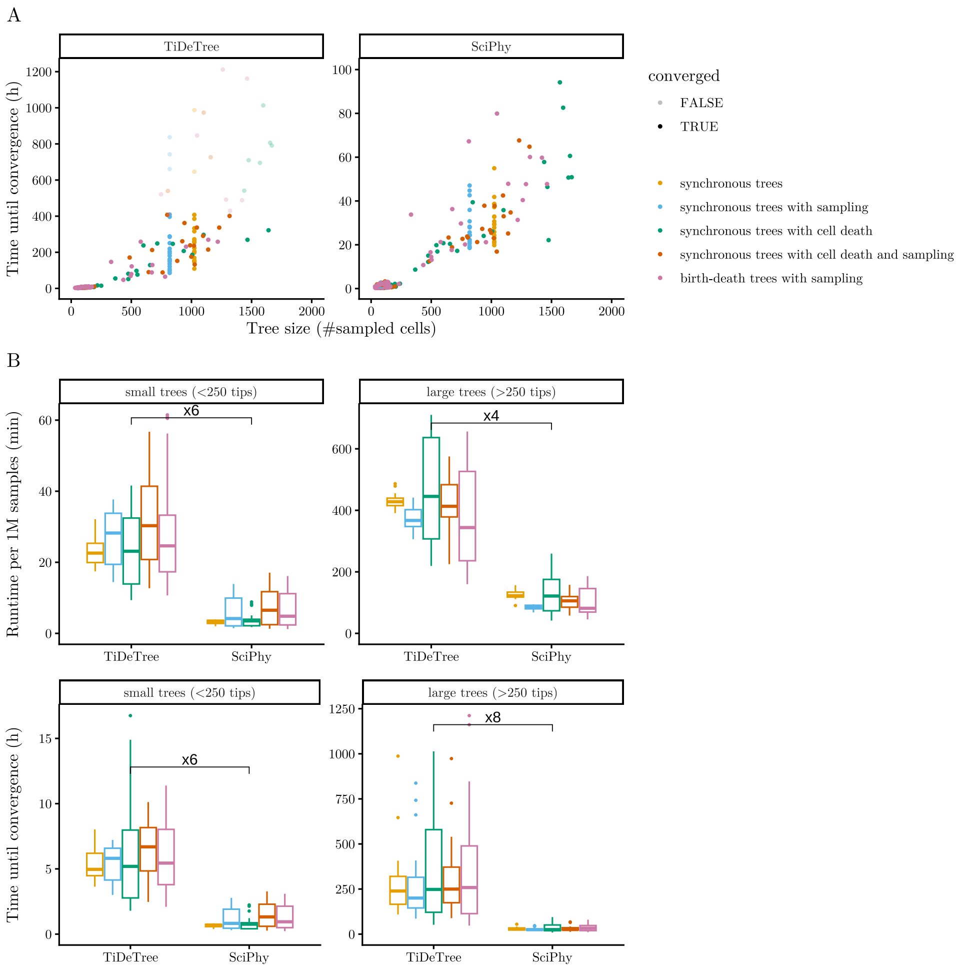


**Fig. S12**: Runtime of Bayesian inference under *TiDeTree* and *SciPhy*. Panel (A) shows the time required for convergence (i.e. MCMC reaching posterior ESS $> 200$, $y$-axis) as a function of tree size ($x$-axis). Panel (B) compares the methods in terms of runtime per million MCMC samples (top) and total time until convergence (bottom). Taken into account are MCMC runs on recordings simulated along the 100 large trees, 60 downsampled trees, and 100 filtered trees per recorder type. Colors indicate the different underlying tree generating processes.

Prospectively, several strategies may reduce the inference runtime. First, running multiple independent chains in parallel and aggregating them for downstream analysis would reduce the wall-clock time required for individual recordings. Second, employing alternative tree and parameter operators – and adjusting their weights –

could enhance mixing, thereby facilitating a more efficient exploration of the state space. Third, the caching mechanism, as already implemented in *SciPhy*, could also be employed in *TiDeTree*. Fourth, integrating the methods with BEAGLE [6], which exploits hardware capabilities of CPUs and GPUs, could speed-up the phylogenetic likelihood calculations. Finally, recent advancements in the Bayesian phylogenetics field offer promising avenues for scaling up the methods to state-of-the-art single-cell lineage tracing datasets. These include the development of targeted operators within the BEAST2 framework [7], which have demonstrated up to a tenfold improvement in mixing and enabled analysis on an empirical dataset with 10,000 sequences. Another significant innovation is Delphy [8], a Bayesian approach that utilizes explicit mutation-annotated trees (eMATs) to streamline likelihood evaluations. Representing all mutation events along branches instead of integrating them out as in Felsenstein's pruning algorithm, massively reduces the computational burden and enables faster MCMC steps, facilitating the analysis of large datasets.

Note that coupling multi-type phylodynamic models to the CRISPR editing models in the Bayesian framework results in even more computationally intensive inference. While analytical solutions exist for the phylogenetic likelihoods under *TiDeTree* and *SciPhy* and the phylodynamic likelihood under the single-type birth-death model [9, 10], the phylodynamic likelihood under the multi-type birth-death model [11–13] needs to be solved numerically. This calculation is also performed at each MCMC iteration – it involves solving a system of ordinary differential equations (ODEs) for every branch, a task that scales with tree size. Furthermore, the phylodynamic parameter space grows quadratically with the number of types, considering all possible transitions among them. Consequently, multi-type inference on larger trees is currently computationally intractable, and we limited our simulations to smaller samples. As algorithms evolve to handle more cells and more complex differentiation landscapes, we anticipate that Bayesian phylodynamics will become a powerful tool for analysing single-cell lineage tracing datasets.

### S6 Filtering out noisy data

As mentioned in Section 2.5, filtering out cells with noisy or missing barcodes induced non-random sampling of lineages in the phylogenies. In Fig. S13, we visualize this effect and compare the inferred and true branch length distributions across population-dynamic models.

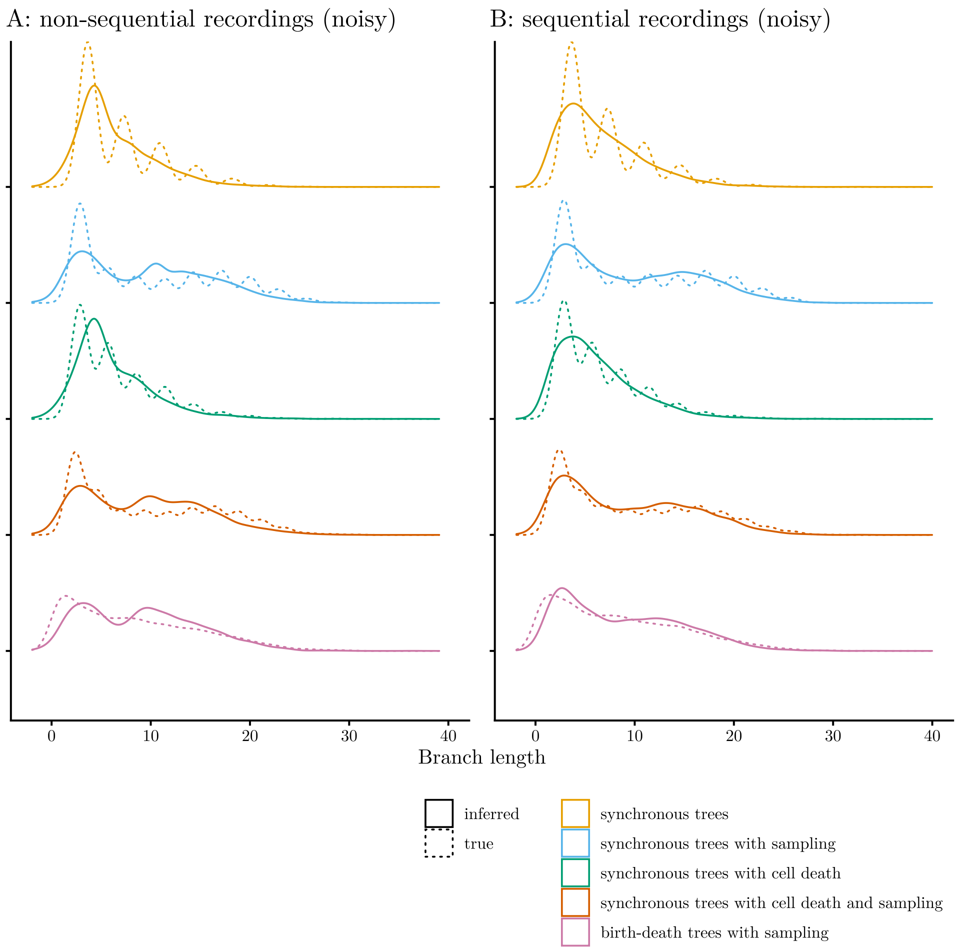


**Fig. S13**: Inferred versus true branch length distributions from filtered (A) non-sequential and (B) sequential CRISPR lineage recordings. Lines indicate smoothed density estimates for the true (solid) and inferred (dashed) branch length distributions. The branch lengths are aggregated across 20 trees per tree generating process (indicated by colors).

### S7 Inferring cell differentiation dynamics

In Fig. S14, we show example multi-type birth-death trees for the two cell type transition scenarios and their inferred counterparts. Note that the posterior and summary trees are 'colored' – the stochastic mapping algorithm [13] was used to place ancestral cell type transitions along the lineages (cf. Sections 2.6 and Methods). Fig. S15 depicts the trees with inflated transition rate estimates. Further, Fig. S16 shows the recovery of transition statistics across simulations. Finally, Fig. S17 summarizes the performance of parameter inference from non-sequential and sequential recordings.

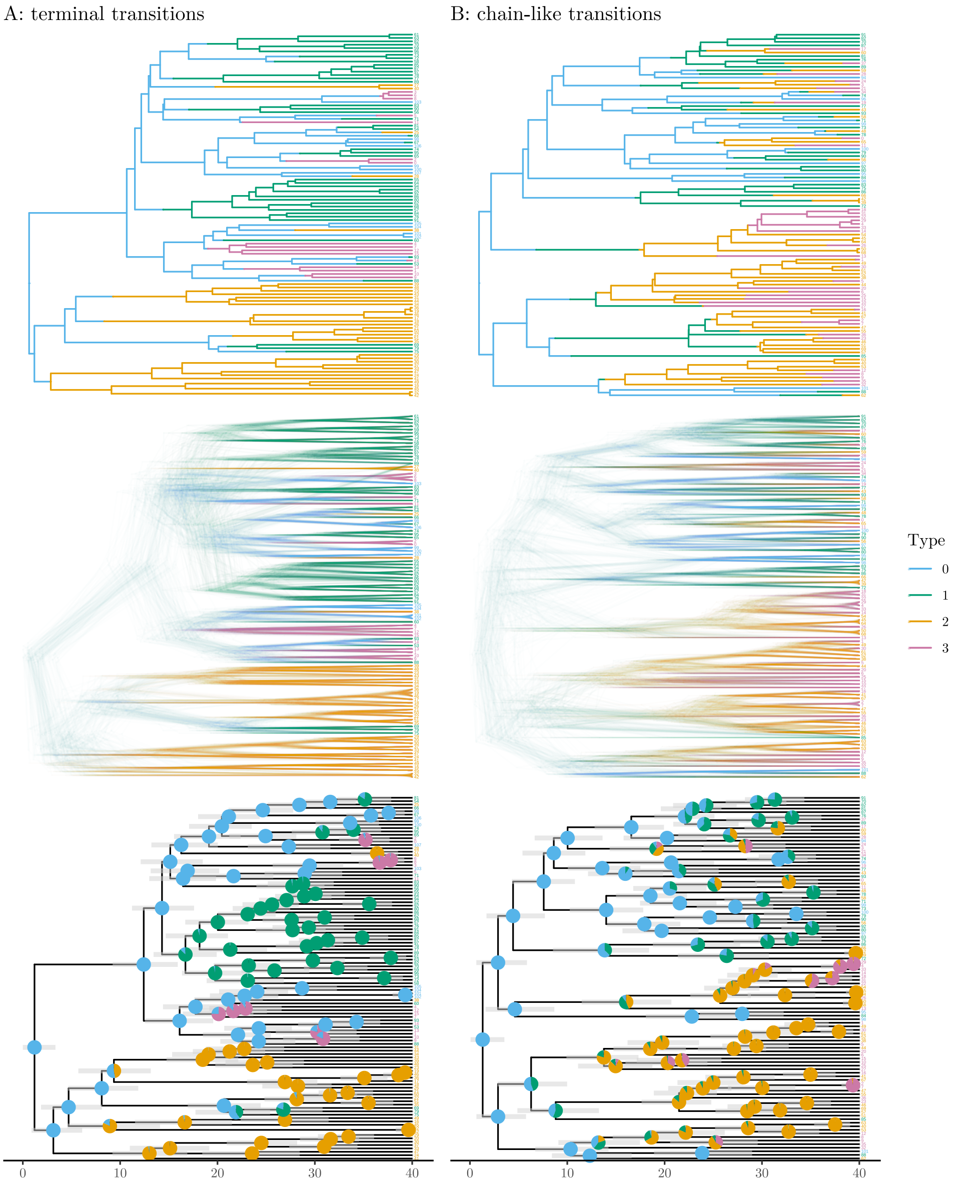


**Fig. S14**: Example multi-type birth-death trees with (A) terminal and (B) chain-like transitions. The panels show from top to bottom: the true tree, a subset of the posterior tree set, the maximum clade credibility (MCC) tree inferred from the corresponding simulated sequential recording. The branches and tip labels are colored by cell type. In the MCC tree, light grey bars indicate the 95% HPD intervals around internal node heights. Pie charts at internal nodes indicate the posterior distribution of ancestral cell types. The trees are time-scaled.

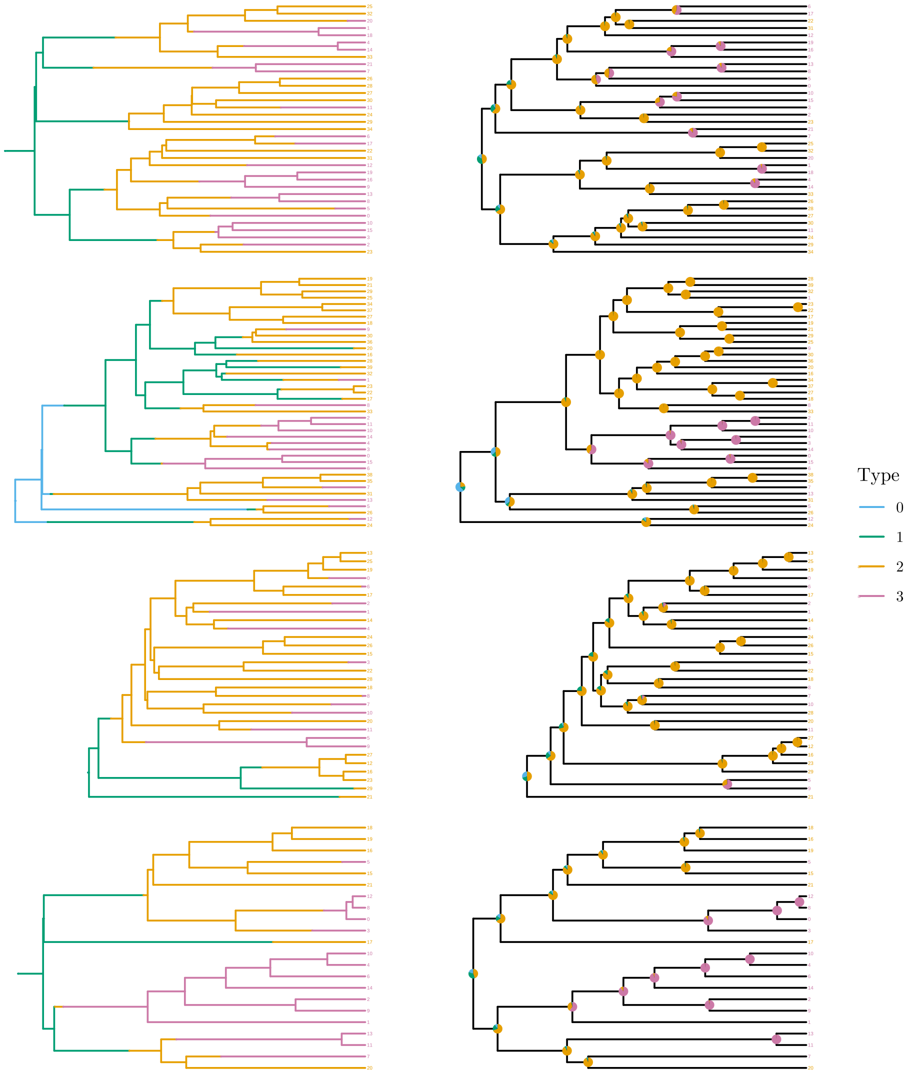


**Fig. S15**: Multi-type birth-death trees with chain-like transitions, for which the transition rates to type 2 were substantially overestimated. The true trees are shown on the left, the corresponding maximum clade credibility (MCC) trees with ancestral cell type probabilities on the right. The branches and tip labels are colored by cell type.

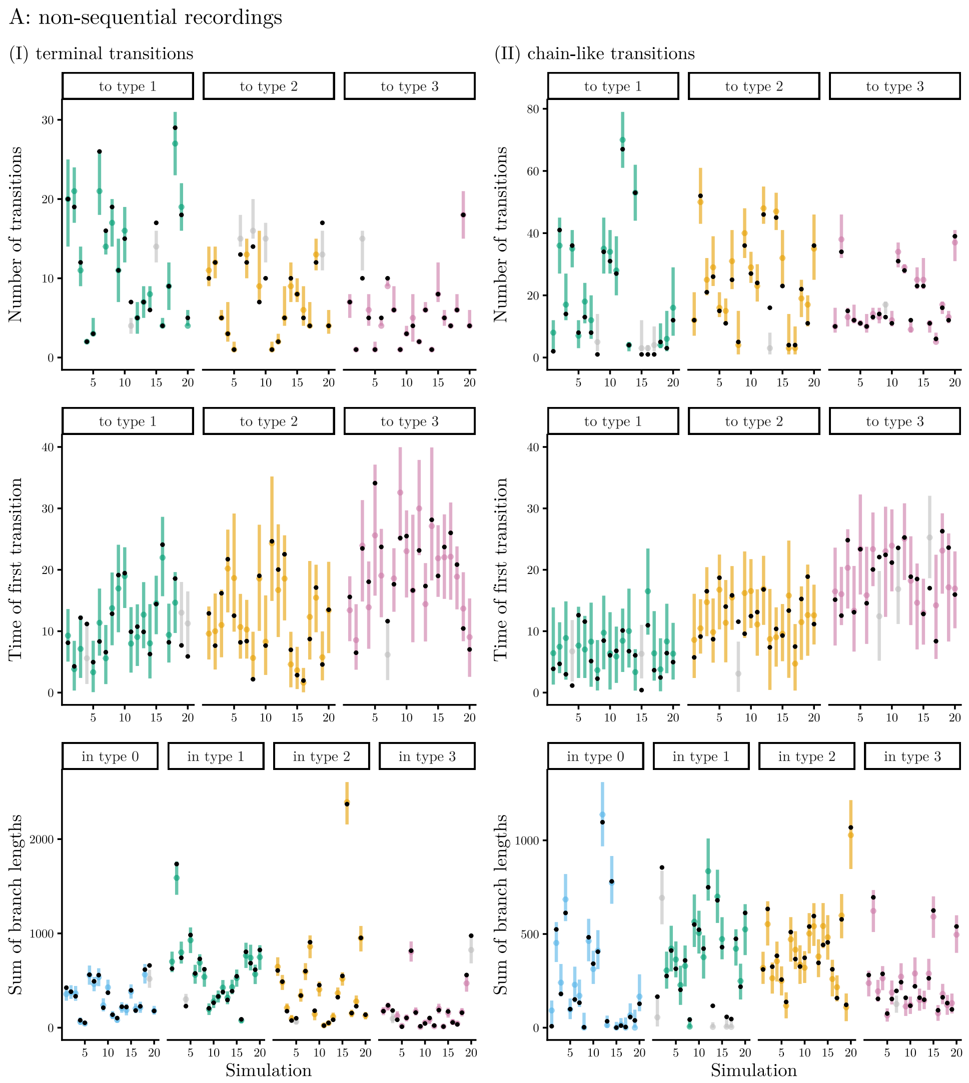


**Fig. S16**: Statistics of type transitions inferred from (A) non-sequential and (B) sequential CRISPR lineage recordings simulated along multi-type trees with (I) terminal and (II) chain-like transitions. In each graph, colored points indicate the medians and bars show the 95% HPD intervals of each statistic (y-axis) per simulation (x-axis), calculated for the posterior tree set. Colors and facets indicate the types. Black points indicate the true values, grey bars indicate simulations, for which the true statistic was not recovered.

B: sequential recordings

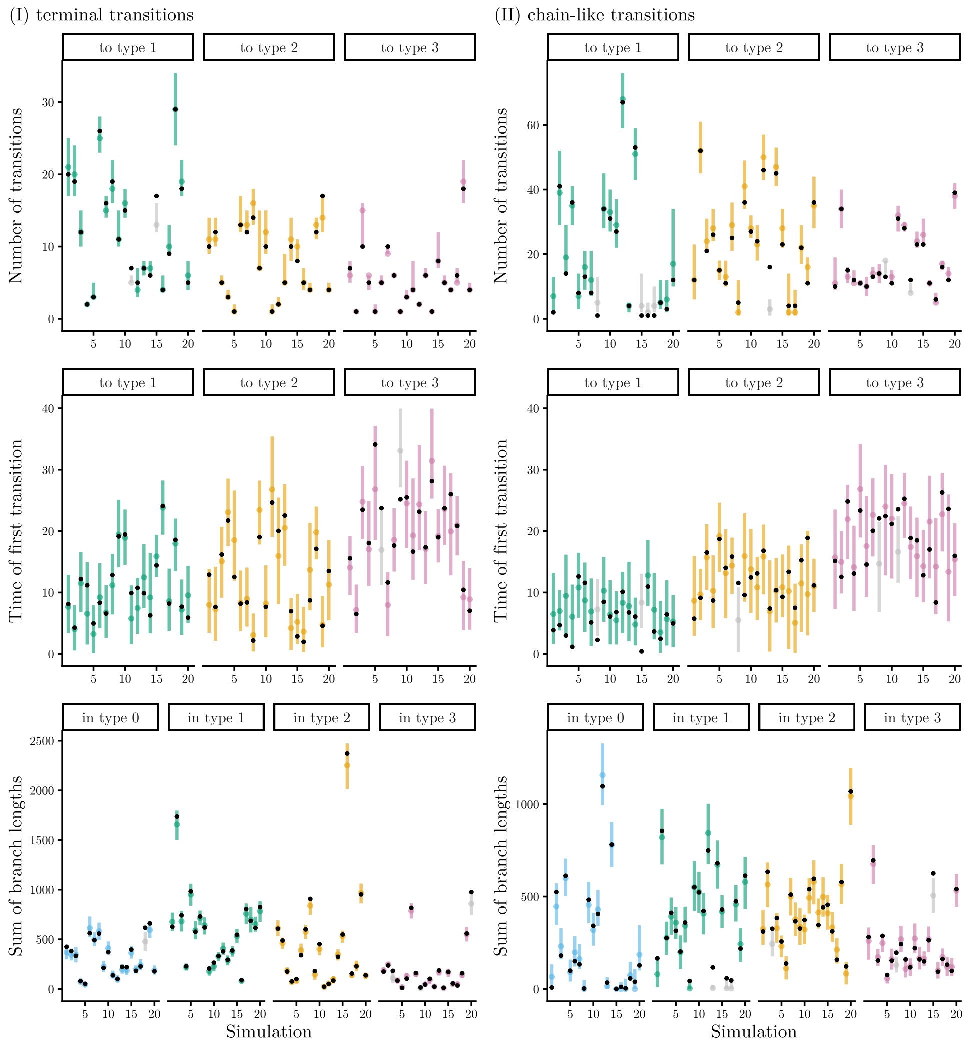


**Fig. S16**: Statistics of type transitions inferred from (A) non-sequential and (B) sequential CRISPR lineage recordings simulated along multi-type trees with (I) terminal and (II) chain-like transitions. (cont.)

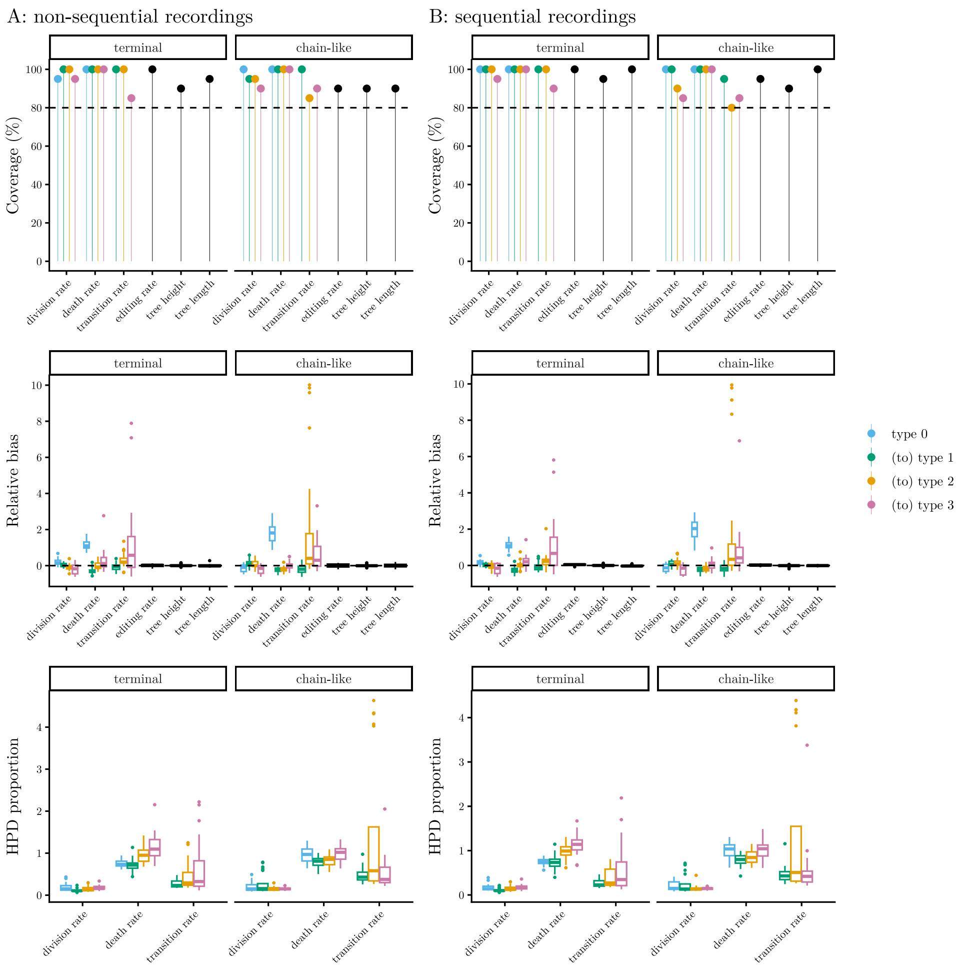


**Fig. S17**: Inference performance on (A) non-sequential and (B) sequential CRISPR lineage recordings simulated along multi-type birth-death trees. The graphs show from top to bottom: the coverage, relative bias and HPD proportion of the inferred parameters. Colors indicate for which cell type the parameters hold. The inference results are summarized across 20 simulations per cell type transition dynamics. Dashed lines display thresholds for comparability.

[7] Bouckaert RR, Weidemüller PH, Gomez LRE, Müller NF. Improving the Scalability of Bayesian Phylodynamic Inference through Efficient MCMC Proposals.

bioRxiv. 2025;p. 2025–06. https://doi.org/10.1101/2025.06.18.660471.
